# Genetic and stochastic basis of phenotypic discordance in 16p11.2 mouse model deletion

**DOI:** 10.64898/2026.07.29.741187

**Authors:** Julian Taranda, Jana Velíšková, Sukalp Muzumdar, Kith Pradhan, Kannan Umadevi Venkataraju, Ramesh Palaniswamy, Rhonda Drewes, Jianjun Sun, Nicolas Dross, Sevin Turcan, Jesse Gillis, Libor Velíšek, Pavel Osten

## Abstract

Recurrent copy number variations such as the 16p11.2 deletion (del/+) represent a significant genetic risk for neurodevelopmental disorders, showing incomplete penetrance and variable expressivity. Here we describe a striking discordance in a genetic mouse model of human 16p11.2 deletion. Using electrocorticography experiments, we detected that approximately 50% of the isogenic del/+ mice exhibited increased susceptibility to seizures from low doses of the convulsant drug pentylenetetrazole (PTZ). Some of these mice also displayed spontaneous epileptiform brain activity. Next, we used two-photon microscopy tomography (STPT) with c-Fos-GFP (c-Fos+) as a reporter of neuronal activity to obtain images of the entire mouse brain. We examined del/+ mice that exhibited low or high responses to PTZ. Low responders exhibited minimal c-Fos+ activation. However, they had activating circuits that suggest the presence of compensatory mechanisms that suppress seizures. These circuits include the reticular nucleus of the thalamus, the pretectal region, the pedunculopontine nucleus, and the superior colliculus. In contrast, high-responder mice exhibited high c-Fos+ expression, corresponding to a fast seizure phenotype. Using whole-brain imaging, we analyzed brain volume and detected significant differences in the cortex between low-responder and high-responder del/+ mice. In the group with higher seizure susceptibility, we observed a notable decrease in cortical volume with significant differences in several brain regions compared to low responders. Additionally, some high-responder mice showed null expansion areas compared to low-responder mice. During the longitudinal behavioral analysis, the high-responder group of mice exhibited more severe sleep disturbances and an increase in repetitive grooming. Finally, to identify transcriptional correlates of the low- versus high-response phenotypes in adulthood, we performed bulk RNA-seq on somatosensory cortex punches from adult del/+ mice stratified by seizure response. Seizure-prone del/+ mice exhibited reduced activity-dependent immediate early gene signatures and coordinated changes in regulatory programs, including decreased expression of transcriptional regulators and metabotropic glutamate receptor genes, compared to low responders. Together, these findings define two distinct adult del/+ subgroups with convergent circuit, structural, behavioral, and molecular signatures.

## Introduction

Human genetic studies have uncovered many risk loci for neurodevelopmental disorders and congenital anomalies, including large copy number variations (CNVs) spanning multiple candidate genes.^1–3^ However, it has been difficult to progress from genetic findings to mechanisms underlying disease, perhaps due to the difficulty in modeling incomplete phenotypic penetrance and genetic complexity of human CNV disorders in genetic mouse models.^4^ One such example is the del/+, which has been associated with a wide range of human developmental and neurological phenotypes, including autism spectrum disorder (ASD), coordination disorders, language disorders, lower mean IQ scores, and various electroencephalogram (EEG) abnormalities and seizures.^2,5,6^ Notably, as with other CNVs, del/+ patients exhibit highly variable phenotypes even among closely related individuals, such as a multiplex family in which the father and proband son share features of autism, intellectual disability, and congenital malformations, but a younger brother carrying the same deletion is largely unaffected.^7^ Such findings are consistent with four mechanisms of origin contributing to the phenotypic inconsistency: 1) variable genetic background, 2) de novo second-hit mutations, 3) environmental factors, and 4) inherent stochastic processes regulating the phenotypic penetrance of CNVs, such as epigenetic variation or stochastic developmental or transcriptional processes. Our study design differs from previous comparisons of del/+ mutant with wild-type (wt) mice, which primarily aimed to detect differences averaged across all mutant mice. These previous studies identified hyperactivity, reduced sleep, subtle abnormalities in cognitive tests and anatomical changes including reduced cortex and caudoputamen and enlarged hypothalamus and midbrain in the wt versus mutant cohort, but failed to identify discordant phenotypes within the mutant mouse cohort.^8–13^ Here, to specifically investigate possible divergent phenotypes in the mouse model of the human 16p11.2 deletion^3^ and to discriminate between the sources of such phenotypic variability, we designed a series of experiments using isogenic del/+ mice maintained in the same environment and with sufficient replicates to exclude the effects of rare de novo mutations. We performed sensitive and unbiased whole brain analyses, coupled with measures of seizures, cortical activity and homecage behavior (undisturbed mice), to identify discordant phenotypes that may be relevant to phenotypes associated with the human syndrome. Next, to detect a possible early genetic bifurcation in del/+ mice, we performed a whole-genome expression analysis on P2 and adult mice to identify the potential genetic cause of this discordant phenotype.

## Materials and Methods

### Animals

Mice postnatal day between 8-12 weeks males and females were used as indicated in the text and figures. Pure 129Sv 16p11.2 del/+ mouse model (backcrossed > 10 generations in 129Sv) provided from Alea Mills lab^3^ were crossed with pure C57BL/6N c-Fos-GFP mouse model (backcrossed > 10 generations in C57BL/6N from the original B6DBA-Tg Fos-tTA, Fos-EGFP & Tg tetO-lacZ, tTA, Jax lab # 008344, resulting in a loss of the second transgene Tg tetO-lacZ, tTA). The *c-fos-GFP* transgene includes all 4 exons and all introns of the *c-fos* gene^14^. The validation of c-Fos^+^ expression with respect to native c-Fos expression was carried out by us preciously.^15^ Experimental mice were F1 generation of background 50:50 C57BL/6N:129Sv from breeding of 16p11.2 del/+ heterozygous male and c-Fos^+^ heterozygous females. The mice were kept on a 12:12 light/dark cycle and given food and water *ad* libitum. All animal procedures were approved by the Cold Spring Harbor Laboratory Animal Care and Use Committee.

## Results

### Del/+ mice show increased vulnerability to seizures and abnormal cortical activity

Given the frequent and variable occurrence of EEG abnormalities and seizures in the 16p11.2 deletion patient population (∼50% and ∼30% of del/+ carriers, respectively),^2^ we first asked whether such changes may be revealed in del/+ mice spontaneously and/or after a dose-dependent exposure to a chemoconvulsant drug PTZ. Seizure activity can be reported using scores for severity, ranging from freezing (1) to myoclonic seizure (jerk/twitch) (2) to clonic seizure (forelimb clonus) (3) to tonic-clonic seizure (4). As shown in (Fig. 1), the del/+ mice exhibited an increased propensity to seizures compared to wt mice as follows. Subthreshold doses of 20 and 30 mg/kg PTZ that caused only freezing in adult wt male mice (*n* = 10 and 12, respectively) evoked a range of behavioral seizures in adult del/+ male mice, suggesting increased seizure propensity in at least some mutant animals: PTZ 20 mg/kg evoked myoclonic seizures in 2 out of 10 mutant mice, and PTZ 30 mg/kg evoked myoclonic seizures in 3 and clonic seizures in 7 out 14 mutant mice. Suprathreshold PTZ dose of 40 mg/kg induced clonic seizures in both wt and mutant mice (Fig. 1A), although the overall onset of the seizure was faster and the seizure duration longer in del/+ mice (Supplementary Fig. 1A and B; all mice were isogenic F1 background 50:50 C57BL/6N:129Sv). A similar increase in propensity to subthreshold PTZ was also observed in female del/+ mice F1 C57BL/6N:129Sv background and in male del/+ mice in 100% C57BL/6 background, indicating that this phenotype is not dependent on sex or mouse genetic background (Supplementary Fig. 1C).

**Figure 1.**
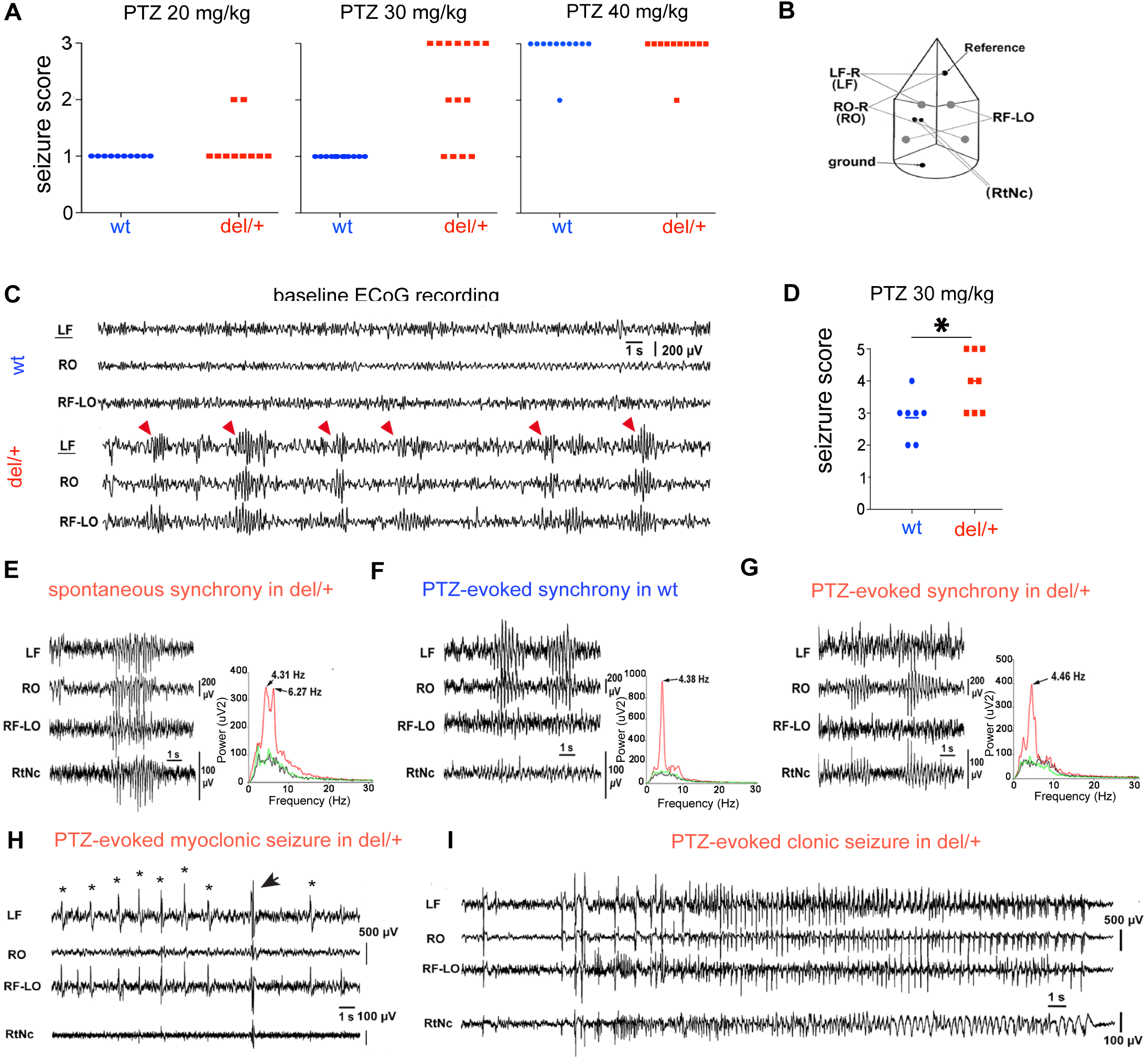
Seizure susceptibility and abnormal cortical activity in del/+ mice. **(A)** del/+ (red) mice show increased PTZ-evoked seizure response: 2 out of 10 mutant mice showed myoclonic seizures after 20 mg/kg PTZ, and 3 and 7 out of 14 mutant mice showed myoclonic and clonic seizures, respectively, after 30 mg/kg PTZ (*p*=0.0002; Fisher’s Exact Test). In contrast, all wt (blue) mice responded only with freezing at these doses. At 40 mg/kg PTZ, 10 out of 11 mutant and wt mice responded with clonic seizures. (**B)** Schema of ECoG electrodes: LF-R = left frontal vs. reference, RO-R = right occipital vs. reference, RF-LO = right frontal vs. left occipital, RtNc = bipolar in the left thalamic reticular nucleus. **(C)** Example of ECoG recording over 30 s illustrates spontaneous synchronous waxing-waning waves of cortical ECoG activity that frequently occurred over a 24-hour recording period (red arrow heads) in 2 out of 8 mutant mice, in contrast to 0 out of 7 wt mice. **(D)** PTZ-evoked scores of ECoG recorded mice. ECoG activity and occurrence of the following: 1 = isolated waxing-waning ECoG waves without freezing, 2 = generalized waxing-waning ECoG waves with freezing, 3 = ECoG spike without a myoclonic twitch, 4 = ECoG spike-and-wave with a myoclonic twitch, 5 = fast poly-spike ictal ECoG activity with a clonic seizure of forelimb. See panels 1H and 1I for ECoG recording examples. Statistical comparison by Mann-Whitney test (\**p* < 0.05). **(E-G)** Analysis of E spontaneous and F-G, 30 mg/kg PTZ-evoked rhythmic activity in F del/+ and G wt mice. Each panel shows (left) 10-sec trace of ECoG recoding and (right) power spectral analyses of baseline activity (black line), waxing-waning ECoG waves (red line), and return to baseline activity after waxing-waning waves (green line). Note the prominent peaks at ∼4.3 Hz in spontaneous ECoG activity in mutant mice E, and PTZ-evoked activity in both genotypes F-G suggesting that the abnormal spontaneous synchrony reflects subthreshold epileptiform activity. In addition, a faster peak at 6.27 Hz in spontaneous activity E suggests a recruitment of a more complex circuitry in the mutant mice. **(H)** At 30 mg/kg PTZ-evoked myoclonic seizure in del/+ mice: spike and wave discharges (marked by “*”) not associated with body twitches (ECoG score 3) were followed by a discharge associated with a body twitch (a myoclonic seizure; marked by an arrow; ECoG score 4); such activity was seen in 5 out of 8 del/+ mice, but only 1 out 7 wt mice. **(I)** At 30 mg/kg PTZ-evoked clonic forelimb seizure in del/+ mice: the seizure started with a few spaced spike-and-wave discharges, followed by fast poly-spikes of decreasing amplitude and frequency (ECoG score 5). This occurred in 3 out of 8 del/+ mice, but 0 out of 7 wt littermates.

To probe these findings further, we employed ECoG recordings to obtain direct measurements of both spontaneous and PTZ-evoked cortical and thalamic neuronal activity (Fig. 1B-I). Recording of spontaneous ECoG activity over a 24-hour period identified episodes of waxing-waning waves synchronized in the cortex and reticular thalamus. These waves indicated abnormal, generalized, spontaneous, rhythmic activity in the cortex and reticular thalamus of 2 out of 8 mutant mice. No such activity was observed in 7 wt littermates (Fig. 1C). Next, in agreement with the initial behavioral observations, increased and varied seizure responses to the 30 mg/kg PTZ dose in del/+ mice were revealed by evaluation of both ECoG recordings and behavioral scoring: PTZ evoked waxing-waning waves synchronized in the cortex as well as in the reticular thalamic nucleus (generalized rhythmic PTZ-induced activity) associated with freezing in 8 out of 8 mutant mice, followed by myoclonic seizures in 6 out of 8, and clonic seizures in 3 out of 8 mutant mice (Fig. 1D and I). In contrast, while all wt mice also responded with generalized rhythmic PTZ-induced activity (synchronized in the cortex) associated with freezing, only 1 out of 7 wt mice showed myoclonic seizure and no wt mice exhibited clonic seizures (Fig. 1D-I). Furthermore, the two mutant mice with spontaneous abnormal synchronous activity (Figures 1C and D) were among those three responding with PTZ-evoked clonic seizures, suggesting that the spontaneous waxing-waning waves occurring in del/+ mice predict increased seizure susceptibility. The similarity between the spontaneous and PTZ-evoked waxing-waning activity is further supported by power spectral analysis of these events, which revealed a common ∼4.4 Hz peak in all three conditions—spontaneous and PTZ-evoked waxing-waning waves in del/+ mice and PTZ-evoked activity in wt littermates (Fig. 1E-G and Supplementary Fig. 1D and E).

### Whole-brain c-Fos mapping reveals two distinct groups in del/+ mice

Our results show that a subset del/+ mice exhibit abnormal synchronous activity and a higher propensity to PTZ-evoked seizures. The variable seizure response may reflect variable phenotypic penetrance, where some mice are more affected than others carrying the same mutation. Alternatively, these results could indicate that del/+ mice are similarly affected but exhibit high phenotypic variability. To differentiate between these two scenarios, we asked whether del/+ mice with a high seizure propensity could be distinguished from del/+ mice with a low seizure propensity by functional and structural brain changes. Such findings would support the existence of two distinct populations del/+ mice with discordant phenotypes. To analyze the underlying whole-brain functional and structural changes, we used our high-resolution STPT imaging assay^15–18^ to 1) map brainwide neuronal excitation, represented by the induction of the immediate early gene *c-fos,*^19,20^ and 2) measure regional volumes of the imaged brains based on the segmentation by the Allen Mouse Brain Atlas (version ARA 2008).^21,22^

In the first set of c-Fos-mapping experiments, we evaluated the distribution c-Fos^+^ cell distribution at undisturbed animals as a proxy for resting brain activity in del/+ and wt littermate mice (homecage group) and i.p. saline solution. This analysis revealed comparable distributions of c-Fos^+^ cells, suggesting an absence of significant differences in resting brain activity between del/+ and wt littermates. Among those injected with saline solution, del/+ and wt mice exhibited similar c-Fos^+^ activation, though some areas were activated exclusively in del/+ mice. (Supplementary Fig. 2, Supplementary Table 1). Next, to search for circuit differences underlying the differences in seizure propensity in del/+ mice, we performed c-Fos^+^-mapping experiments after exposure to 30 mg/kg PTZ, using the induction of c-Fos^+^ as a cellular reporter of PTZ-evoked brain activation.^19,20^ Based on our initial observations, we analyzed the data by dividing the del/+ mice into two phenotypic groups: 1) “low responder” (LR) mutant mice exhibiting freezing behavior and 2) “high responder” (HR) mutant mice exhibiting clonic seizures. Both groups were compared del/+ mice injected with saline solution (saline control group; Fig. 2A and B, Supplementary Table 2A and B). As expected, c-Fos^+^ activation evoked by 30 mg/kg PTZ in the LR and HR mice was markedly different due to their different seizure phenotypes: freezing behavior in LR mutant mice was associated with modest, mainly subcortical activation, whereas clonic seizure in HR mutant mice was associated with broad brain wide activation that included the entire cortex, hippocampus, caudoputamen, many midline thalamic nuclei and other subcortical areas (compare Fig. 2A and B; and see Supplementary Tables 2A, B and E). Surprisingly though, a closer analysis of the modest LR brain activation pattern revealed a number subcortical areas that were activated only in the LR mutant mice and not in the HR mice and also not in wt mice with comparable PTZ-induced freezing and clonic seizures (Fig. 2 C and D; Supplementary Table 2 C and D) (the LR-specific areas are marked by arrows in Fig. 2A-D; note that wt mice were injected with a higher dose of 35 mg/kg PTZ in order to achieve comparable PTZ-evoked responses as del/+ mice, which is further quantified in Supplementary Fig. 3). Notably, among the LR-specific structures many belonged to circuitries capable of seizure controlling and terminating activity,^23^ including the thalamic reticular nucleus and parafascicular nucleus,^24–26^ midbrain reticular nucleus,^27^ superior colliculus,^28,29^ ventral tegmental nucleus,^27,30^ several midbrain and hindbrain raphe nuclei^27^ and several structures of the ventromedial medulla^31^ (Figures 2A and E, Supplementary Table 2). Furthermore, the pedunculopontine nucleus, which together with the superior colliculus is a key part of the basal ganglia seizure terminating network,^29,32^ was also much more prominently activated in the LR compared to HR mutant mice (Fig. 2A, B and E). Taken together, these findings point to compensatory brain circuit mechanisms acting to suppress and terminate seizures selectively in the LR mutant mice and not in the HR mutant mice.

**Figure 2.**
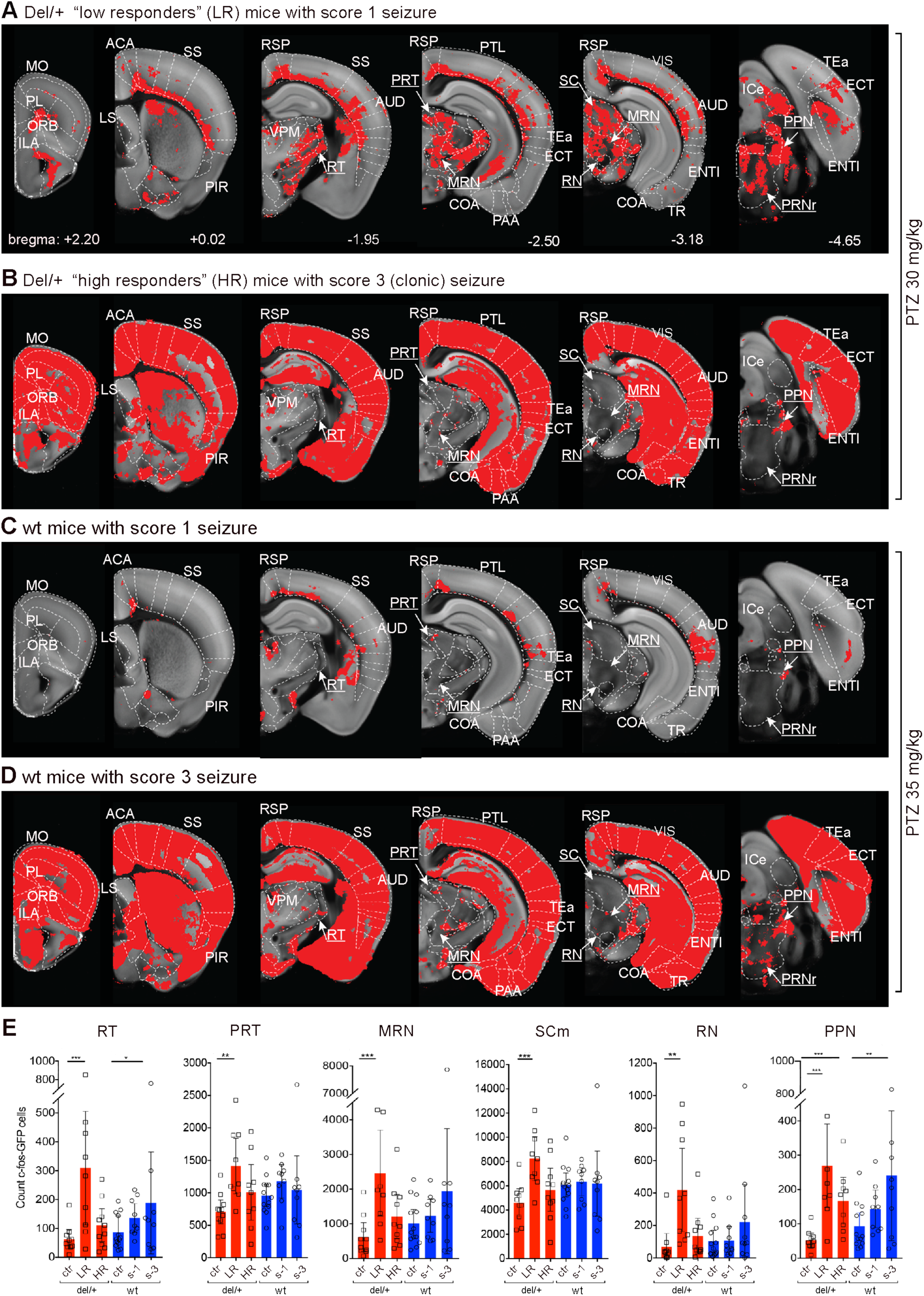
Quantification of PTZ-evoked seizures by whole brain c-Fos^+^ mapping detect in del/+ LR group, specific anticonvulsant circuit stimulation. **(A-D)** Statistical comparison of c-Fos^+^ cell distribution between control mice injected with saline and mice injected with PTZ, with significant increases representing higher brain activation evoked by PTZ shown as red areas overlaid on a grayscale RSTP mouse brain^15^. **(A)** PTZ-evoked freezing behavior corresponded to mainly subcortical activation in the LR del/+ mice. **(B)** PTZ-evoked clonic seizures induced broad activation across the cortex, hippocampus, caudoputamen and midline thalamic nuclei in the HR del/+ mice. Notably though, despite much strong activation in the HR mice, a number of areas known to act as anticonvulsant brain circuits^26–28,60^ were activated only in the LR mice, including the reticular thalamic nucleus (RT), midbrain reticular nucleus (MRN), superior colliculus (SC), red nucleus (RN), pontine reticular nucleus (PRNr), and the pretectal region (PRT), while the pedunculopontine nucleus (PPN) that acts together with SC was also activated more robustly in LR than HR mice (the names of these structures are underlined in A-D panels and the regions are highlighted by arrow; other abbreviations are listed in Supplementary Table 2) (saline injected *n* = 10, LR *n* = 9, HR *n* = 10 mice). **(C)** PTZ-evoked freezing in wt mice, modest cortical and subcortical activation lacking any of the seizure controlling brain structures seen in LR del/+ mice. **(D)** PTZ-evoked clonic seizures in wt mice induced broad activation comparable to the del/+ HR response. Note that del/+ mice were injected with 30 mg/kg PTZ whereas wt mice with 35 mg/kg PTZ to normalize for lower seizure propensity in wt mice (wt saline injected *n* = 13, wt score 1 *n* = 9, wt score 3 *n* = 9 mice). The statistical *q* value for voxel-based analysis was *q*<0.05. **E.** Bar graph representation of c-Fos^+^ cell numbers in selected brain areas with known anticonvulsant activity and highlighted above (A-B) (ctr = saline injected) (del/+, red) and (wt, blue). The *q* values were derived by negative binomial regression corrected for multiple comparisons by FDR; \**q*<0.05, \*\**q*<0.01, \*\*\**q*<0.001.

### Quantification of brain volume shows structural differences between del/+ LR and HR mice

Given the clear and robust differences in PTZ-evoked brain activity between the LR and HR mutant mice, we next search for structural differences that may be related to the different functional phenotypes. To this end, we quantified brain volumes in the imaged brains from the mice used in the experiments described above. First, we carried out a two-cohort comparison between naïve del/+ mice and wt littermates, pooling the brains from the homecage and saline injected groups to form a baseline mutant versus wt comparison (Fig. 3A). This experiment confirmed and extended on previous findings from MRI-based studies using the traditional two-cohort comparison,^3,10^ demonstrating that the mutant mice differed from their wt littermates by 1) volume reductions of the caudoputamen, hippocampus, and several cortical areas, including the frontal, motor, somatosensory, auditory and visual cortex, and 2) volume enlargements of the hypothalamus, superior colliculus, and periaqueductal gray (Fig. 3A, Supplementary Table 3A). Second, we compared separately the LR and HR mutant mice to wt littermates with PTZ-evoked freezing and clonic seizures, respectively. This analysis revealed two important and novel findings: The LR mutant brains showed subcortical expansions seen in the earlier two-cohort comparison, including the periaqueductal gray and superior colliculus, but only minor reductions in the cortex. In contrast, the HR mutant mice showed large brain volume reductions, including all cortical areas, caudoputamen, hippocampus and many thalamic nuclei, but without any subcortical enlargements (Fig. 3B and D and Supplementary Table 3B-D). These data suggest that the previously described phenotype of combined brain volume reductions and expansions in del/+ mice is not entirely accurate. ^3,10^ The data are better explained by the existence of two del/+ groups with divergent phenotypes.

**Figure 3.**
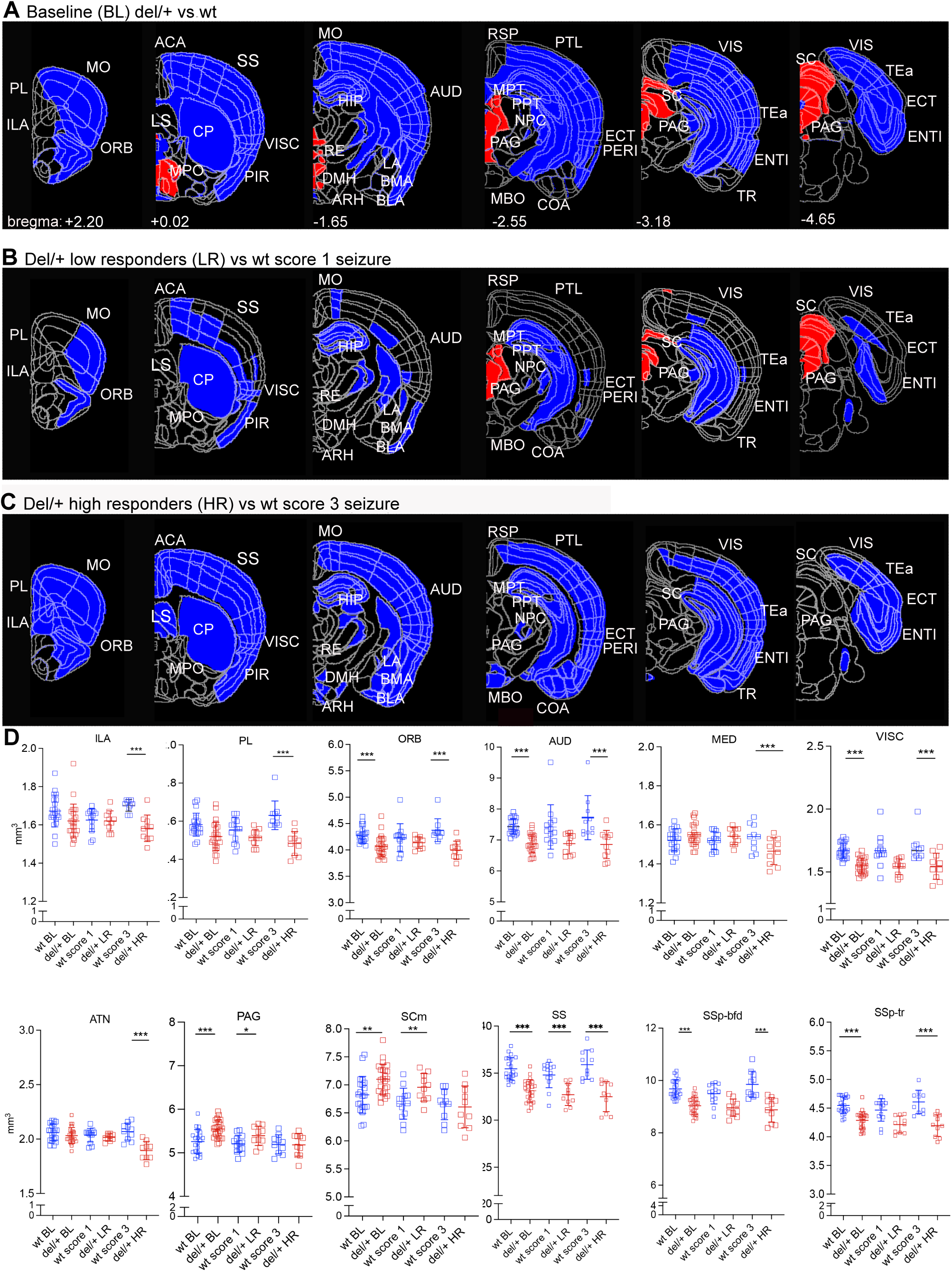
Quantification of brain volumes shows structural differences between del/+ LR and HR mice. Volume analysis of STPT imaged mouse brains^15,61^ with larger areas shown in red and smaller areas in blue. **(A)** Two-cohort comparison between del/+ and wt mice baseline (BL) groups. In agreement with previous MRI. studies^3,10^, del/+ brains showed smaller cortical areas, including motor (MO), orbital (ORB), somatosensory (SS), visceral (VISC), piriform (PIR), auditory (AUD), visual (VIS), parietal (PTL), ectorhinal (ECT) and entorhinal (ENT) cortex, as well as caudoputamen (CP), and hippocampus (HIP) that contrasted with larger hypothalamus (medial preoptic area, MPO, dorsomedial nucleus, DMH), periaqueductal gray (PAG), and superior colliculus (SC) (Supplementary Table 3) (del/+ *n* = 24, wt *n* = 21). **(B)** Comparison between del/+ LR and wt score 1 mice with PTZ-evoked freezing response revealed only modest volume reductions of cortex but persistent increases of PAG. and SC in the LR mice (LR *n* = 10, wt score 1 *n* = 13). **(C)** Comparison between del/+ HR and wt score 3 mice with seizures revealed prominent volume reductions the cortex, caudoputamen hippocampus, no volume increases in del/+ HR mice (HR *n* = 10, wt score 3 *n* = 10). Additional volumes reduction don’t seen in the baseline Fig. 1A, was detected comparison included prelimbic (PL), infralimbic (ILA), anterior cingulate (ACA), and retrosplenial (RSP) cortex, as well as subcortical latera; septum (LS), lateral, basomedial and basolateral amygdala (LA, BMA, BLA), and cortical amygdala (COA). The statistical comparisons were done by negative binomial regression corrected for multiple comparisons by FDR, red for expansion \**q*<0.05, blue for reduction \*\*\**q*<0.001. (**D)** Bar graph shows the volume of selected brain areas from the three comparisons above. Each point corresponds to one animal. Del/+ (red), wt (blue). Primary somatosensory, barrel field (SSp-bfd), and primary somatosensory, trunk (SSp-tr). The statistical *q* values were derived by negative binomial regression corrected for multiple comparisons by FDR; \**q*<0.05, \*\**q*<0.01, \*\*\**q*<0.001.

### The HR group showed a more pronounced change in the behaviour of the mice compared to the LR group

If the CNV effect can lead to stable differences in brain anatomy and seizure susceptibility, we next asked whether the LR and HR mutant del/+ mice may also exhibit divergent behavioral phenotypes related to developmental disorders? To address this question, we performed a series of behavioral analyses followed by PTZ experiments to post-hoc identify the LR versus HR mutant mice. First, we video-monitored homecage behavior of 17 mutant mice and 12 wt littermates using the homecage behavior (CleverSys Inc) in 3 sessions over 1 week. Second, after collecting all behavioral data, we injected each mouse three-times with 30 mg/kg PTZ, done once per week for three weeks, to test the stability of the seizure phenotype over time in individual animals. As shown in Fig. 4A, the repeated PTZ treatment again divided the del/+ mice into two LR and HR phenotypic groups, as 9 out of 17 mutant mice showed freezing (LR = score 1) in all three tests and the remaining 8 out of 17 mutant mice showed myoclonic or clonic seizures (HR = score 2 or 3) in all three tests (Fig. 4A). In contrast, the same treatment induced freezing (score 1) in 11 out 12 wt mice in all 3 tests (Fig. 4A). The analysis of the preceding homecage behavior revealed the following results: When all del/+ mice were pooled together and compared to wt mice in a two-cohort analysis, the mutant mice showed previously described hyperactivity (Fig. 4B and C).^3,10^ However, when we subdivided the del/+ mice into LR and HR mice, the LR del/+ mice showed hyperactivity (Fig. 4F and G), but the HR del/+ mice also showed increased stereotyped repetitive behavior in the form of increased self-grooming and decreased sleep (Fig. 4H and I, and Supplementary Fig. 4A and B).

**Figure 4.**
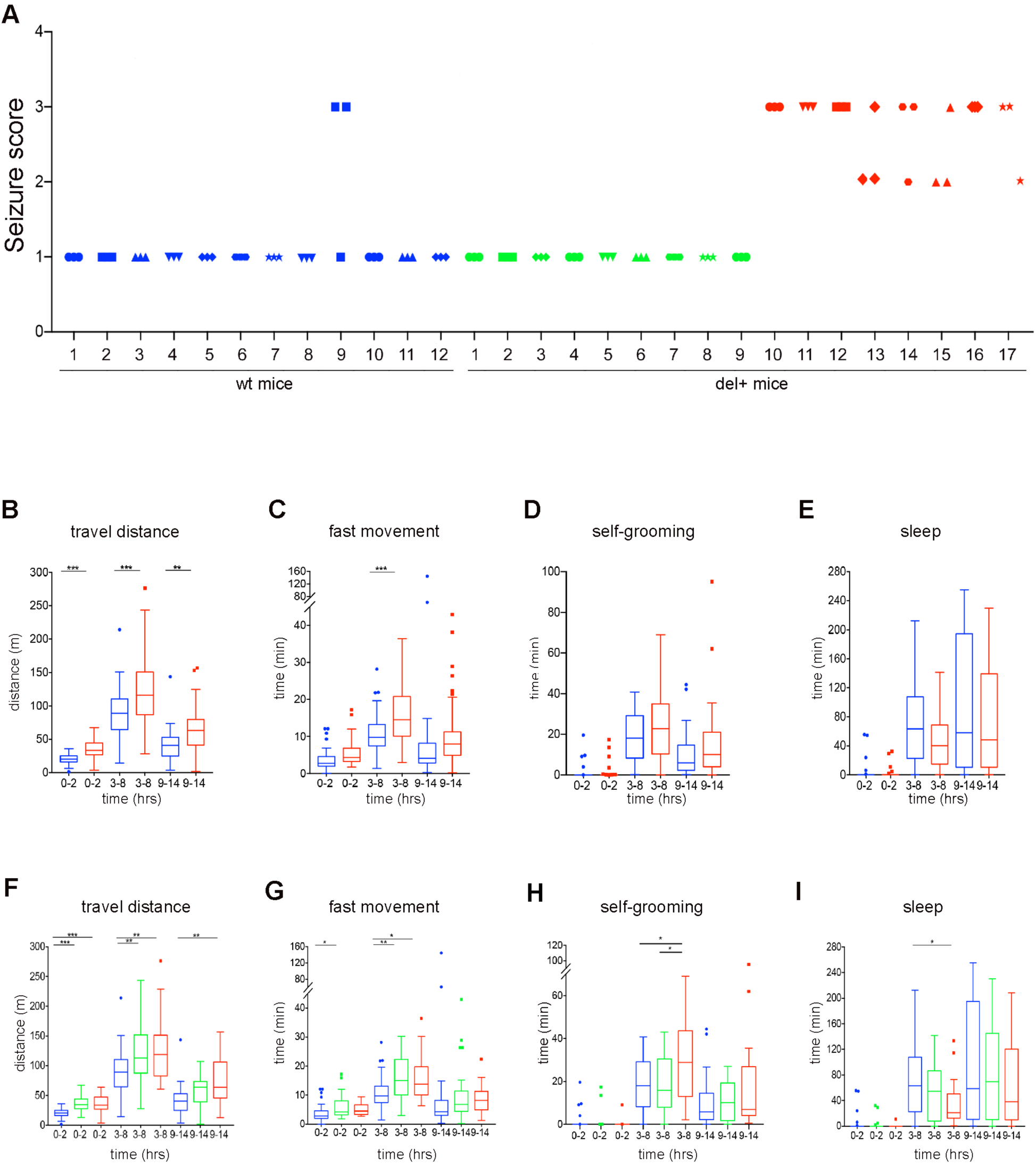
Quantification of behavioral changes in del/+ LR and HR mice. **(A)** Repeated PTZ (30 mg/kg) injections (once per week over 3 weeks) were done after behavioral testing showed in panels **(B-I)**. The response in del/+ mice confirmed the existence of two phenotypic LR (green) and HR (red) groups: 9 out 17 mutant mice showed freezing “low response” to all 3 PTZ injections, whereas 8 out of 17 mutant mice showed myoclonic and clonic seizures “high response” to all 3 PTZ injections. In contrast, 11 out 12 wt (blue) mice always showed freezing. **(B-E)** Two-cohort comparison between all del/+ (red) mice (*n* = 17) and wt (green) littermates (*n* = 12) identified hyperactivity expressed as increased travel distance and fast-moving time (B-C) in del/+ mice. **(F-I)** Comparison between del/+ LR (green) (*n* = 9) and HR (red) (*n* = 8) mice selected based on their seizure response and wt mice (*n*=12) revealed hyperactivity in both LR and HR mutant mice (F-G) but increased self-grooming and reduced sleep only in the HR mutant mice (H-I). For B-E, statistical comparisons were done using one-way ANOVA with Sidak′s comparisons test to compare two groups. Travel distance (0-2 *p*<0.001; 3-8 *p*<0.001; 9-14 *p*<0.01), fast movement (3-8 *p*<0.001). For F-I, statistical comparison were done using one-way ANOVA and Tukey multiple comparisons test with wt, LR and HR responder mice to compare three groups in F-I. Travel distance (0-2 wt-LR *p*<0.001; 0-2 wt-HR *p*<0.001; 3-8 wt-LR *p*<0.01; 3-8 wt-HR *p*<0.01; 9-14 wt-HR *p*<0.01), fast movement (0-2 wt-LR *p*<0.05; 3-8 wt-LR *p*<0.01; 3-8 wt-HR *p*<0.01), self-grooming (3-8 wt-HR *p*<0.05; 3-8 LR-HR *p*<0.05) and sleep (3-8 wt-HR *p*<0.05).

### RNA isolation reveals robust adult gene signatures in del/+ mouse

Previous results demonstrate that del/+ mice can be divided into two groups based on phenotypic severity. We then performed a genome-wide transcriptional study using bulk RNA purified from the SS of del/+ mice at two different ages to analyze gene expression changes associated with discordant phenotypes. The SS was selected because it showed a significant reduction in volume between LR and HR, provide easy access and minimized discomfort for animals undergoing biopsies for subsequent seizure testing (Fig. 3D). Group 1 consisted of male pups at P2, which corresponds to 216 days post-conception in humans.^33^ During mid-late fetal and early postnatal life, genes within the 16p11.2 deletion are highly co-expressed in the human brain.^34^ Group 2, consisting of undisturbed adult male mice (homecage), provided a gene expression profile in the SS between 2-3 months of age (Fig. 5A). First, we analyzed the expression of the genes involved in the deletion compared to their diploid control at both ages. For genes within the del/+ region, we observed a uniform reduction in gene expression in the right hemisphere (dorsal pallium) for P2 and SS in adult brains and their blood samples (approximately 50% reduction) (Supplementary Fig.5A and B). However, genes within the deletion region were more effective at distinguishing del/+ genotypes in brain samples than in blood samples (Supplementary Fig. 5C and D). When genes in the deletion region were excluded, principal component analysis at P2 did not separate del/+ and wt genotypes in both brain and blood (Fig. 5B). In adult mouse brain samples, transcriptional variance allowed for the separation of the two genotypes into two significant clusters, with the majority of del/+ mice falling into in cluster 2 (6 of 10 mice). However, such clear separation was not evident in the blood samples (Fig. 5C and Supplementary Fig. 5E and Supplementary Table 4). Differential expression analysis revealed minimal differences between the brain and blood samples when the genes in the deletion were excluded. Only a few genes were differentially expressed (Supplementary Fig. 5F and G). Therefore, we decided to use the top 100 upregulated and downregulated genes as possible signatures to facilitate genotype separation (referred to as signatures in this paper) and to analyze an aggregate genes effect in del/+ mice. The brain and blood upregulated and downregulated gene signatures efficiently discriminated del/+ mice from wt mice at both ages (Fig. 5D-E). Next, we applied the P2 brain up and downregulated signatures to the adult RNA and vice versa to determine whether any of these signatures could separate the genotypes. The most robust signature detected was 100 top-upregulated genes in the adult brain, which also differentiated P2 pups (Fig. 5F, Supplementary Table 5). Surprisingly, the P2 and adult blood samples lacked the resolution to separate the two genotypes in another age group (Supplementary Fig. 6A and B). When we analyzed the common genes in the signatures between P2 and adult brain mice, we noted that most of these shared genes in both age groups were involved in cell metabolism such as *Lnpp5f, Ralpbp1, Anhgef25, Senp2, and Coq10b* (Fig. 5G). Among the top 10 upregulated genes in P2, we detected differential expression of genes involved in brain development in the isocortex, such as *MAST1* and *Kirrel3*,^35,36^ in the top 10 genes in adults, we identified a gene related to cell metabolism (*Ttr*)^37^ (Supplementary Fig. 6C and D, Supplementary Table 6). Furthermore, the *Rps13-ps1* pseudogene appears to be differentially expressed in P2 and adults (Fig. 5F and G and Supplementary Fig. 6C and D). We then performed Gene Ontology (GO) enrichment analysis on the signatures. The 100 top upregulated genes in P2 animals showed pathways affecting postsynaptic density, ionotropic glutamate receptor binding, and clathrin-dependent endocytosis. In the upregulated signature of the adult group, GO analysis revealed significant changes in vascular organization, astrocytes, and endothelial cells. The signature of the top 100 downregulated genes showed a significant decrease in the expression levels of genes involved in the ERK pathway and MAP kinase activity (Fig. 5H and Supplementary Table 7).^38^ Next, to analyze possible cell-types involved, we performed a cell-type deconvolution using the upregulated signatures. In the P2 group, this analysis indicated the presence of transcriptional signatures associated with neuronal development, mainly in the deep layers of the dorsal pallium. In the adult group, vascular features (VLMC) and astrocytes (Astro) were the most enriched cell-type signatures. For the downregulated signatures, the adult brain transcriptional alterations were significantly enriched for L4/5 IT. The blood samples were not enriched for isocortex markers in either group^39^ (Fig. 5I).^40,41^

**Figure 5.**
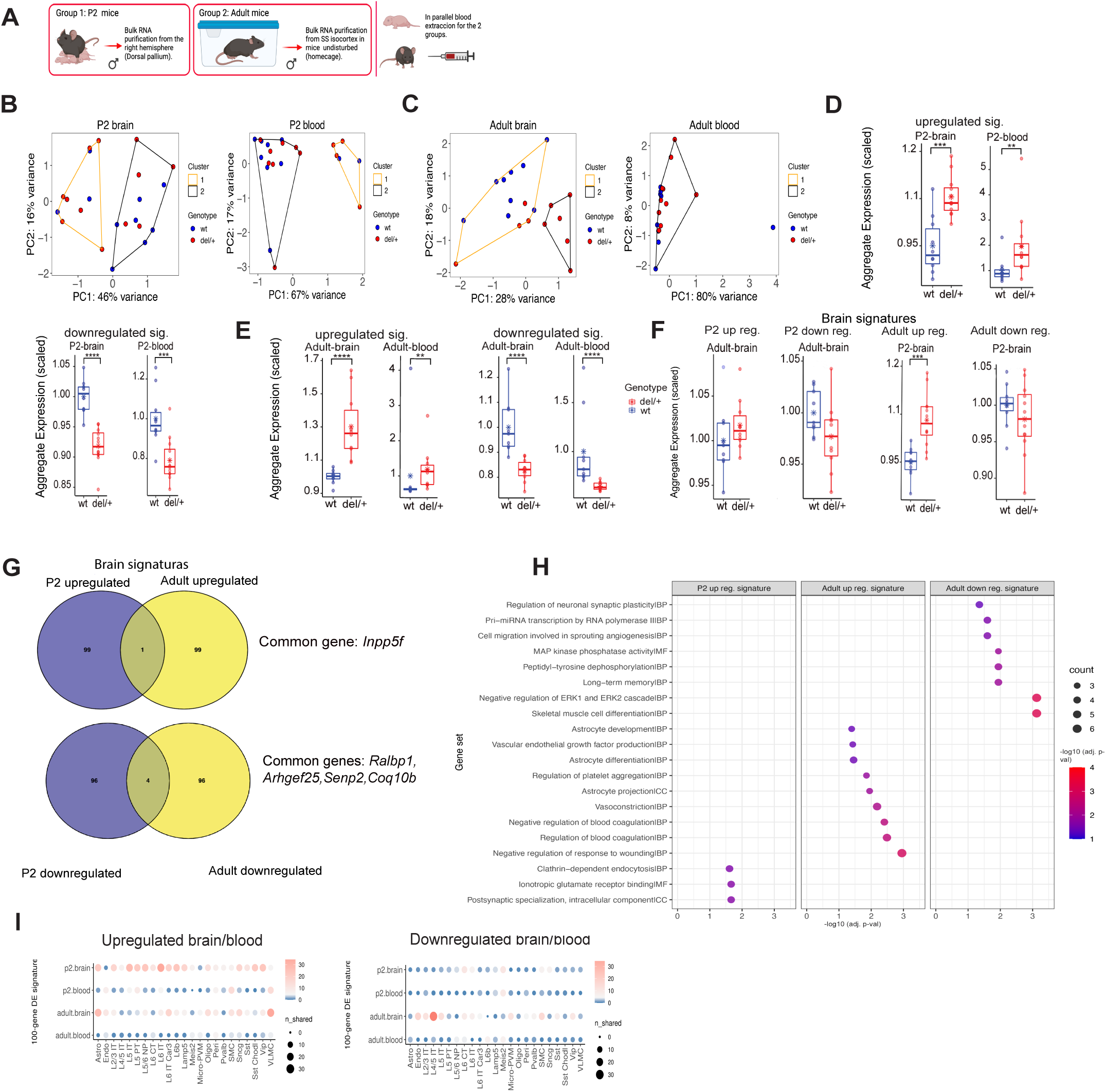
RNA isolation reveals a critical moment in the gene expression profile during P2 and adult in the del/+ mouse. **(A)** Group 1: Bulk RNA purification from P2 littermate pups (wt *n*=10, del/+ *n*=12,). Group 2: Bulk RNA purification from adult littermates (wt *n*=9, del/+ *n*=10). Blood samples were collected for RNA purification for both groups. (**B)** Principal component analysis in wt (blue) and del/+ (red) littermates of P2 mice in brain and blood samples, to identify possible clusters (yellow and black), p values are not significant. (**C)** Principal component analysis in wt (blue) and del/+ (red) littermates in brain tissue and blood of adult mice, *p* value was significant from brain tissue samples (** for *p*<0.010-Fisher’s exact test). (**D)** Top 100 upregulated genes signature from brain RNA of P2 mice in brain and blood (*** for *p*<0.001) (** for *p*<0.01). Top 100 downregulated genes signature from P2 brain RNA in brain and blood (**** for *p*<0.0001) (*** for *p*<0.001). These genes are shown as aggregated expressions in all cases. **(E)** Top 100 upregulated genes signature from brain and blood RNA in adult mice (**** for *p*<0.0001) (** for *p*<0.01). Top 100 downregulated genes signature from brain and blood RNA in adult mice (**** for *p*<0.0001) (**** for *p*<0.0001). (**F)** RNA brain tissue signatures from P2 and adult applied in adult and P2 respectively for genotype differentiation (** for *p*<0.01). (**G)** Venn diagrams of common genes for the up-and downregulated genes signature between P2 and adult. (**H)** Hypergeometric G.O. enrichment analysis in the up-and downregulated genes signature between P2 and adult mice, showing significant ontology pathways with adjusted -log10 p-values (* for *p*<0.05=1.3), (BP- biologically process, CC-cellular compartment, MF-molecular function). (**I)** The top 100 up- and downregulated genes from P2 and adult mice, enrichment analysis hypergeometric cell type marker set analysis (MSEA) -log2 (* for adj *p*<0.05=10).

### Del/+ HR group has a higher number of downregulated transcription factor genes than the del/+ LR group

Finally, to determine the transcriptional signatures associated with discrepancy phenotype in the del/+ mice, we collected cortical biopsies from the SS, and after mice recovery, we injected with PTZ (30 mg/kg i.p.) and classified them as either LR or HR, after blood samples were collected (Fig. 6A). Bioinformatic analysis showed that the expression of genes within the deletion region was reduced by 50 % in del/+ mice. When we separated the del/+ mice based on LR and HR, we did not detect significant differences between the genes in the deletion (Supplementary Fig. 6E-F, Supplementary Fig. 7A-B). When we excluded the genes contained in the deletion region, PCA analysis based on the most variable genes was not able to separate the LR and HR groups (Fig. 6B, Supplementary Fig. 7C-D and Supplementary Table 4). Following the above strategy, we generated signatures with the 100 most up- or downregulated genes in the del/+ compared to the wt mice. For the brain samples, the upregulated and downregulated signatures discriminated the wt and del/+ mice, and the del/+ LR and HR groups compared to wt. However, the blood signatures did not separate the different genotypes (Supplementary Fig. 7E, Supplementary Table 8). When we applied the signature from the PTZ-treated group to the adult group, it failed to discriminate the del/+ from the wt (Supplementary Fig. 7F). Subsequently, we applied the upregulated gene signature from the adult (homecage) to the PTZ-treated group, which was able to separate the del/+ mice from the wt mice. Although this signature could not separate LR and HR mice, the HR group showed a trend towards increased expression compared to the wt and LR groups (Fig. 6D, Supplementary Table 8). The downregulated signature from the adult (homecage) effectively separated the PTZ-treated del/+ from the wt mice. For this signature, del/+ LR mice were similar to the wt animals, whereas the del/+ HR group showed significant differences compared to wt group (Fig. 6E, Supplementary Table 8). Notably, we identified a group of IEGs that were downregulated in this signature (23 IEGs out of 100 genes in the signature), and we then examined a specific IEG signature in the seizure group in more detail.^42^ Here, we found that the del/+ HR mice had significantly lower IEGs expression than the wt group. The del/+ LR and wt showed no significant differences (Fig. 6F, Supplementary Table 8). Interestingly, we found that the adult homecage downregulated signature and the IEGs signature had the greatest overlap compared to the others. (Supplementary Fig. 7G). The blood samples were not efficient to separate the genotypes using the adult homecage signatures and IEGs signature (Supplementary Fig. 8A and B). Then, to identify pathways associated with the LR versus HR seizure phenotypes, we performed differential expression separately for del/+ LR vs wt and del/+ HR vs wt, and derived gene signatures using the same approach as above (top 100 up- and down-regulated genes from each comparison). We next compared pathway enrichments across these two wt-referenced signatures. While no pathways were significantly enriched among genes upregulated in del/+ HR vs wt, several pathways were enriched among genes downregulated in del/+ HR vs wt and were absent or reduced in the del/+ LR vs wt downregulated signature (Fig. 6G; Supplementary Table 7). Notably, the HR-downregulated signature included glutamatergic signaling genes (*Grm2, Grm4*) and a higher number of annotated regulatory genes (including 4 homeobox genes) than the LR-downregulated signature (Fig. 6H; Supplementary Tables 5 and 7). To provide regulatory context for these signatures, we highlight representative regulators within the LR/HR downregulated gene sets (Fig. 6H). The downregulated program in del/+ HR included neuronal identity/developmental regulators (*Foxg1, Pou3f2/Pou3f3, Satb2*) and state regulators (*Nr1d1*, *Foxo3*), whereas the downregulated program in del/+ LR included activity-associated regulators (*Egr3)* together with chromatin and post-transcriptional regulators (*Phc1, Mbnl2, Celf3*). Together with the pathway enrichment results (Fig. 6G; Supplementary Table 7), these observations support the interpretation that LR/HR differences reflect coordinated shifts in gene-regulatory programs rather than isolated single-gene effects.

**Figure 6.**
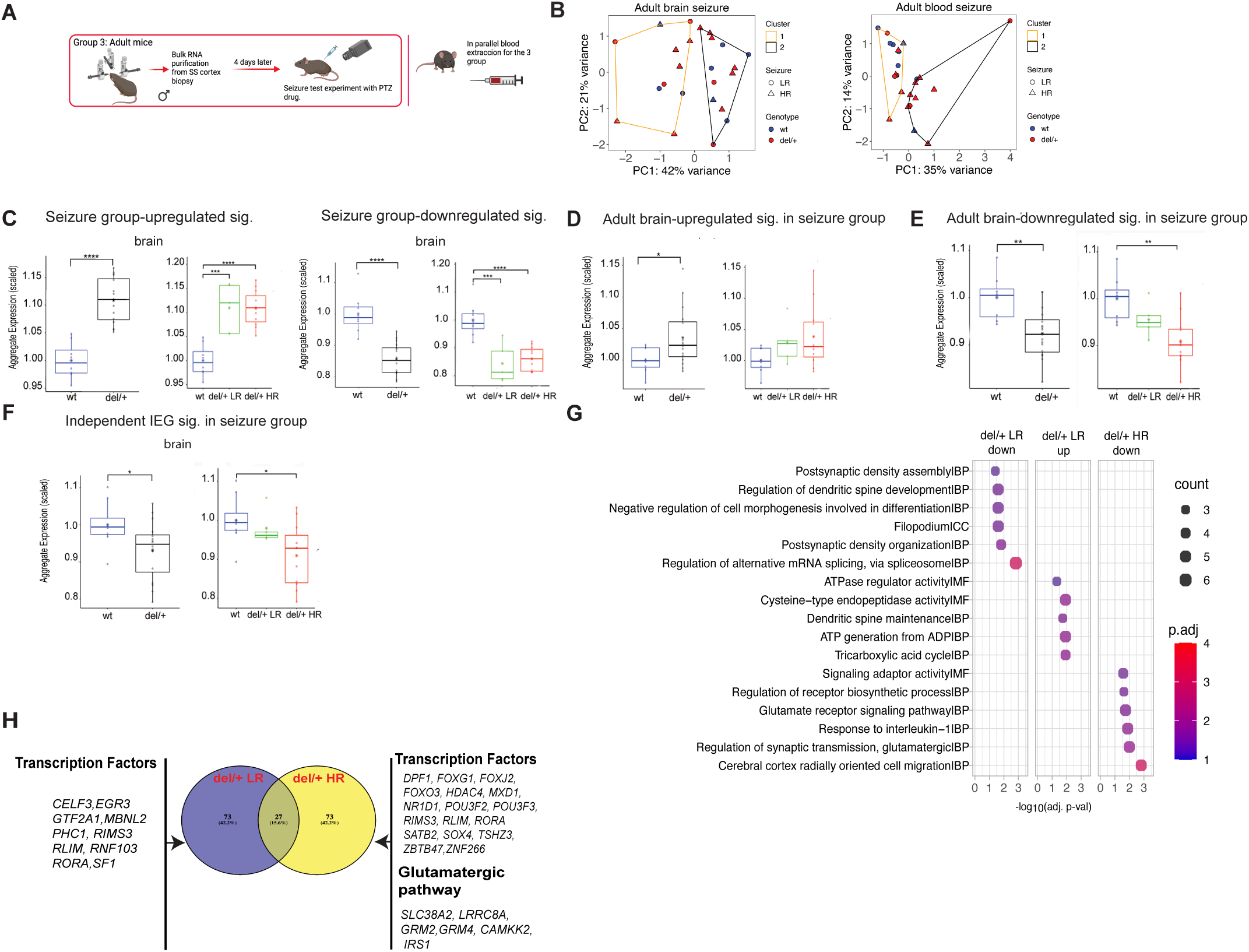
Del/+ HR animals present different genetic profiles than del/+ LR animals. (**A)** Group 3: RNA isolation from the SS cortex biopsies, seizure test and blood extraction. (**B)** Principal component analysis of brain tissue and blood samples in del/+ PTZ-treated adult (del/+ LR, del/+ HR) and wt littermate, *p* values are not significant. (wt *n*=8, del/+ LR *n*=6, del/+ HR *n*=11, littermates). (**C)** Top 100 genes signature upregulated (brain samples- wt vs. del/+ (black) **** for *p*<0.0001, wt (blue)vs. del/+ LR (green)** for *p*<0.01, wt vs. del/+ HR (red) **** for *p*<0.0001), and downregulated ( brain samples- wt vs. del/+ *** for *p*<0.001, wt vs. del/+ LR ** for *p*<0.01, wt vs. del/+ HR *** for *p*<0.001). **(D)** Upregulated signature from RNA adult brain tissue applied in del/+ (black), LR (green), HR (red), and wt (blue) groups. (brain samples-wt vs. del/+ * for *p*<0.05). (**E)** Downregulated signature from RNA adult brain tissue applied in del/+ (black), LR (green), HR (red), and wt (blue) groups (brain samples-wt vs. del/+ ** for *p*<0.01, wt vs. del/+ HR *** for *p*<0.001). **(F)** Immediate-early genes (IEGs) signature applied in the del/+ (black), LR (green), HR (red), and their wt (blue) littermate mice groups (wt vs del/+ * for *p*<0.05, wt vs. del/+ HR * for *p*<0.05). **(G)** Hypergeometric GO enrichment analysis in the up and down-regulated gene signatures in LR and HR groups mice showing the most prominent significant ontology pathways differences -log10 p-values (*p*<0.05=1.3), (BP-biologically process, CC-cellular compartment, MF-molecular function). **(H)** Common genes, and unique genes affected in the seizure group, LR, and HR groups in del/+ mice.

## Discussion

The findings from the 16p11.2 mouse model provide valuable insights to help interpret the phenotypic variability observed in patients with the same deletion. First, since we used isogenic F1 mice and observed phenotypic divergence even among littermates (see Supplementary Fig. 4D, and Supplementary Table 9), the distinct LR and HR phenotypes can be considered comparable to the phenotypic discordances described in monozygotic twin (MZ) studies of CNV syndromes.^43–45^ Second, the consistent emergence of LR and HR phenotypes in a large group of animals (over 70 mutant mice) suggests that stochastic developmental processes, such as epigenetic variability and random fluctuations in lineage commitment and brain wiring likely contribute to divergent neurodevelopmental outcomes.^44,46–48^ This supports the notion that phenotypic discordance in MZ twins, as well as incomplete penetrance and variable expressivity in broader Neurodevelopmental disorders (NDDs) populations, may be driven in part by extragenetic stochastic events, independent of secondary mutations or variable genetic background. Third, abnormal EEG and seizures are frequently observed in autism and other NDDs, but whether this is clinically significant or represents a mere epiphenomenon remains unresolved.^49,50^ Our data in HR group, demonstrate a phenotypic clustering of epileptiform activity, pronounced brain structural changes, and abnormal behaviors, indicating a common mechanism of origin for these deficits. Identifying putative compensatory circuit mechanisms to terminate seizures in the LR mutant mice with normal seizure propensity, only modest brain structural changes, and largely normal behavior further supports a direct phenotypic interaction.

Fourth, RNA-seq data revealed no clear genotype-based clustering in P2 pups (excluding genes within the deletion), likely due to because of the high variability among littermates. In contrast, the adult homecage group exhibited significant cluster separation, with more pronounced genotype-related differences. At P2, most of the differentially expressed genes were related to postsynaptic organization and glutamate receptor binding. This suggests that the P2 stage may precede the full manifestation of genotype-related transcriptional differences, which likely become more pronounced during postnatal development. In rodents, the critical period of synaptogenesis occurs within the first three postnatal weeks.^51,52^ Recent analysis in humans have shown that the genes included in the 16p11.2 deletion are essential in late embryonic and postnatal life^34^. Future RNA analyses should therefore focus on juvenile animals. Previous studies have reported postnatal lethality in a subset of del/+ pups.^3^ In our cohort, 17% of pups died by P1 (data not shown), resulting in a significantly smaller number of del/+ mice. This early loss suggests that more severely affected animals may not have been captured in our datasets. Interestingly, RNA from P2 and adult mice blood samples showed fewer changes than brain tissue data but still reflected gene dosage differences within the deletion. The upregulated gene signature in adults was sufficient to distinguish between the two genotypes in P2. Further transcriptomic analysis in adults revealed that the downregulation of gene signatures included suppression of the MAPK and ERK1/2 pathways. The ERKs play a critical role in corticogenesis by regulating the cell cycle in proliferating neural precursors. It has been reported that genetic ablation of ERKs results in altered brain cytoarchitecture and physiological deficits.^53^ A large body of data demonstrates that ERK1/2 controls multiple stages of brain development and maturation.^54–56^ Interestingly, HR mice exhibited a significant reduction in IEG signatures compared to WT and LR animals. This reduction may stem from the inactivation of ERK1/2, which impairs IEG induction, or from abnormal EEG activity. High seizure susceptibility can blunt IEG expression and desensitize widespread neuronal activation in the brain, which is consistent with accelerated c-Fos responses during seizures.^57^ Notably, HR mice also showed marked downregulation of transcription factors, 20 out of the top 100 downregulated genes were transcriptional regulators. Upon analyzing the downregulated signatures, we discovered that the HR group exhibited striking repression of transcription factors, including four homeobox genes, compared to the LR group. This severe downregulation may be involved in the broader, more significant cortical reduction compared to LR littermates, which affects normal cortical development. It also creates an additive effect in the gene expression profiles downstream of transcription factors.

Finally, the glutamate receptor pathways were affected in the HR group. Specifically, the expression of *Grm2* and *Grm4*, which encode mGluR2 and mGluR4, respectively, was reduced. mGluR2 is mainly presynaptic and controls glutamate in the presynaptic vesicles. Using its antagonist LY34149558^58^ or knock-out mice^59^ increases extracellular glutamate, facilitating the release of glutamate for the presynaptic vesicles. These changes may underlie the heightened PTZ sensitivity in HR mice, in which even low PTZ doses triggered more severe seizure phenotypes. Collectively, our data reveal the emergence of two phenotypically distinct del/+ subgroups, likely representing a stochastic developmental bifurcation. This divergence may result from epigenetic variation, environmental influences, or intrinsic noise in gene regulation during development. These findings have direct implications for understanding variable outcomes in human CNV syndromes, where shared genotypes can result in highly divergent phenotypes.

## Data availability

The bulk RNA raw data is availability in NCBI at GSE212785 ‘‘Genetic and stochastic basis of phenotypic discordance in neurodevelpmental 16p11.2 deletion syndrome.’’ https://www.ncbi.nlm.nih.gov/geo/subs/

## Acknowledgements

We thank Kristin Baldwin (Columbia University) for critical reading and editing the manuscript and Michael Wigler and Ivan Iossifov (CSHL) for helpful discussions and comments. We thank the Histology Core at CSHL and Qing Gao. The Histology core is partially supported by NIH Support Grant 5P30CA045508. We thank the LAR facility at CSHL, especially to Eileen Earl, supervisor of Lab animal resources and to Michael Labarbera. We thank Elena Ghiban and Sara Goodwin as a part of the Next Generation Sequencing Core Facility.

## Funding

This work was supported by grants R01-MH096946 and SFARI 204719 to P.O., and funds from Robertson Research Fund and Stanley Institute for Cognitive Genomics at CSHL to P.O. This work was supported by a grant RC4-NS072966 to L.V., 5R01-NS-092786 to J.V. as well as by Behavioral Phenotyping Core facility at the NYMC. This work was supported by the grant R01MH113005 at CSHL to J.G. This work was supported by the German Cancer Aid, Max Eder Program grant number 70111964 (S.T). This work was supported by DFG grants TU 585/1-1 and 585/1-2 (S.T., J.T.). DFG Mercator fellowship for J.T. This work was supported by CRC1324 by N.D.

## Author Contributions

J.T. and P.O. designed the experiments, analyzed the results, and wrote the paper; J.T. carried out all experiments except ECoG recordings; J.V. contributed to the design of the behavioral experiments. J.V. and L.V. carried out and analyzed ECoG studies, contributed to seizure studies, and commented on the manuscript; K.P. devised and supervised statistical analyses; N.D., K.U.V. devised and supervised STPT data analyses; J.S. performed the brain tissue biopsy for the bulk RNA experiments; S.T., S.M., and J.G. analyzed the data from bulk RNA experiments; R.P., and R.D. were involved in the maintenance of the mouse colony at CSHL.

## Competing interests

The authors report no competing interest in the research of the manuscript.

## Supplementary material

Additional material is available online at Brain. This includes portions of the Materials and Methods section, as well as the Supplementary Figures and Tables.

## Supplemental information

### Supplemental Figures 1-8

**Supplemental** Figure 1 Related with Figure 1

**Supplemental** Figure 2 Related with Figure 2

**Supplemental** Figure 3 Related with Figure 2

**Supplemental** Figure 4 Related with Figure 4

**Supplemental** Figure 5 Related with Figure 5

**Supplemental** Figure 6 Related with Figure 5

**Supplemental** Figure 7 Related with Figure 6

**Supplemental** Figure 8 Related with Figure 6

### Excel Supplementary Tables 1-9

Table 1 Excel file containing additional data too large to fit in a PDF, related to Figure S2

Table 2 Excel file containing additional data too large to fit in a PDF, related to Figure 2

Table 3 Excel file containing additional data too large to fit in a PDF, related to Figure 3

Table 4 Excel file containing additional data too large to fit in a PDF, related to Figure 5/6

Table 5 Excel file containing additional data too large to fit in a PDF, related to Figure 5/6

Table 6 Excel file containing additional data too large to fit in a PDF, related to Supplementary Figure 6C and D

Table 7 Excel file containing additional data too large to fit in a PDF, related to Figure 5/6

Table 8 Excel file containing additional data too large to fit in a PDF, related to Figure 6

Table 9 Excel file containing additional data too large to fit in a PDF, related to Figure 1

## Notes

### Competing Interest Statement

The authors have declared no competing interest.

