## Supplemental Figures for "Genetic and stochastic basis of phenotypic discordance in 16p11.2 mouse model deletion"

### Materials and Methods

**Animal surgeries.** The respective IACUC of Cold Spring Harbor Laboratories and New York Medical College (NYMC) approved all procedures in the animals. Male mice were subjected to surgical implantation of ECoG electrodes using stereotactic frame (Heinrich Kopf, Inc.). Mice were introduced to deep isoflurane anesthesia (5% in $O_2$ for induction in the induction chamber, 2% in $O_2$ for maintenance using an inhalation mask) Depth of anesthesia was monitored by toe-pinch reflex every five minutes. The skull surface was exposed by skin incision. Jeweler's screws were used as a reference and ground electrodes and placed in the nasal bone and behind the lambda, respectively. Silver ball electrodes connected to a connector were used for the cortical recordings and were placed epidurally. The cortical electrodes were positioned symmetrically bilaterally over the sensorimotor (frontal) cortex and over the visual (occipital) cortex. A bipolar wire electrode (Plastics One) was stereotaxically lowered to the thalamic reticular nucleus (20$^{o}$ angle from the sagittal plane; anteroposterior from bregma -0.6 mm, lateral 2.6 mm, depth of 4.0 mm; the mouth bar set at -3.5). All electrodes including the screws were covered with dental acrylic. After the surgery, the animals were placed on a heating pad until fully ambulatory and then returned to their home cage.

**Recordings of brain electrical activity.** After minimum one week of recovery, animals were placed in cylindrical recording containers for 24 hours of habituation. After habituation, the mice were connected to wired preamplifiers (Pinnacle Technology) and recorded (combined electrocorticography (ECoG) and video) for 24 hours for sleep stages. The lights were turned on at 6:00 am (06:00) and turned off at 6:00 pm (18:00). After additional 3 days, the animals were injected with convulsant drug pentylenetetrazole (PTZ) 30 mg/kg intraperitoneally (i.p.) to determine their seizure susceptibility including ECoG response. ECoG evaluation was done by two independent observers blinded to animal identity for the presence of waxing-waning rhythmic waves (spontaneous or PTZ-induced) with the amplitude > two-fold background ECoG activity amplitude, synchronized over cortical recordings (= generalized rhythmic ECoG activity) and for presence of PTZ-elicited spike-and-waves or ictal activity (polyspike-and-wave). For further analysis, Sirenia Seizure Pro (Pinnacle Technology) was used. Specifically, power spectra analysis and seizure discovery using "line length" measures were employed.

**Seizure experiments in 16p11.2 del/+ mice.** 16p11.2 del/c-Fos$^{+}$ and c-Fos$^{+}$ littermates (C57Bl/6N/129sve F1 hybrids) male mice were group-housed before the tests. Five days before experiments, adult (8-12 weeks old mice) were transferred to a designated area in the animal room and separated single per cage (with environment enrichment) to lower the variability in baseline c-Fos$^{+}$ expression[1] (note that the 5-day isolation with enrichment is too short to evoke behavioral and neuroendocrine stress responses in adults).[2] All experiments were done between 10:30 am and 12:30 pm and the mice were sacrificed between 1:30 pm and 3:30 pm. Each animal was taken from its cage, injected i.p. with saline solution or PTZ in the biological safety cabinet class II, returned in its cage with a digital camera from the side to analyze the evoked seizure response. Three groups of animals were analyzed: mice in the homecage without any handling or perturbation, control mice injected with saline solution and mice injected with different concentrations of PTZ The animals were video-recorded to quantify the behavioral score of seizure and after 3 hours they were killed by transcardial perfusion with saline and 4% P.F.A. and the brains were prepared to serial two-photon tomography (STPT) imaging[3] (see below).

**Pentylenetetrazole (PTZ).** PTZ is a chemical convulsant frequently used in seizures studies, acting at least partly by blocking the GABAA receptor.[4] PTZ (SIGMA, P6500-25G) was dissolved in 0.9% of saline solution and used 20, 30, 35 mg/kg for different experiments.

**Quantification of PTZ-evoked responses.** The behavior was video-recorded and the initial 30 min were manually scored off-line by two independent observers to quantify and classify the seizure-related behavior: Score 0 – normal behavioral without freezing; Score 1 – freezing = sudden behavioral arrests and immobility; Score 2 – myoclonic seizure = neck jerking, twitching and tailing, whole body twitch; Score 3 – clonic seizure = repeated clonus of one or both forelimbs (sometimes also head) while animal keeps upright position. The behavior was video-recorded, and the initial 30 min were manually scored off-line by two independent observers blinded to animal identity and PTZ injection to quantify and classify the seizure-related behavior. Scoring system has been adopted to reflect primarily generalized PTZ-induced seizures[5] (Score 0 – normal behavioral without freezing; Score 1 – freezing = sudden behavioral arrests and immobility; Score 2 – myoclonic seizure = neck jerking, twitching and tailing, whole body twitch; Score 3 – clonic seizure = repeated bilateral forelimb clonus (sometimes also head) while animal keeps upright position. In experiments described in Fig. 1A three mice with no response to PTZ were excluded from the analysis: one wt and one del/+ mouse at 20 mg/kg of PTZ, and one del/+ at 30 mg/kg of PTZ. The latency of PTZ-evoked response was measured as time to onset the first behavioral manifestation. The recovery time was defined as the time post behavioral seizure manifestation when the mouse crossed the cage-length and was walking with lifting all four paws from the ground.[6]

**Histology and STPT imaging.** Mice were perfused transcardially with ice-cold 0.9 % saline solution followed by 4% paraformaldehyde (P.F.A.) (diluted in 0.2 M phosphate buffer) for 7 min at 7 ml/min, at 3 hrs post behavioral experiment. Brains were post-fixed in 4% P.F.A. for 24 hours at 4 °C, then transferred to 0.1 M glycine solution (diluted 0.1 M phosphate buffer) for 48 hrs at 4

°C, then stored in 0.1 M phosphate buffer at 4°C until imaged. Imaging was done as previously described.[3,7] In brief, brains were embedded in 4% agarose in 0.05M PB, cross-linked in 0.2% sodium borohydrate solution (in 0.05 M sodium borate buffer, pH 9.0-9.5). The brains were imaged with TissueCyte 1000 (Tissue Vision) 2-photon microscope with integrated vibratome at 1µm x-y resolution and Z spacing of 50 µm. This generated 280 serial sections datasets. The raw images tiles (16-bit tiff) were corrected for illumination, stitched in 2D open-STP procedure.[3] 2-photon excitation wavelength was 910 nm; 560 nm dichroic mirror (Chroma, T560LPXR) and bandpass filters (Semrock FF01-520/35 an) were used to separate green signal and red background channel.

**Automated cFos-GFP (c-Fos$^+$) cell counting.** c-Fos$^+$ neurons were automatically detected by a convolutional network (C.N.) (ID: 164) trained to recognize nuclear neuronal cell body labeling with F-score 0.89.[3] c-Fos$^+$ induction in the cerebellum was not analyzed in the current study, because of a high a false positive rate in this brain region. For each dataset the brightness of the signal of each sample was normalized by the mean and standard deviation of tissue auto-fluorescence signal from a coronal section at a bregma position of +1.0 mm.

**3D brain registration.** 3D registration methods were the same as described by us previously, at 20 µm × 20 µm × 50 µm pixel size using affine and b-spline transformation in Elastix[8], an image registration toolbox based on Insight's ITK. The precision of the registration was measured by the displacement of 13 landmark points in 6 different mouse brains after warping each dataset onto the average RSTP (reference serial two-photons tomography) brain.[3]

**Quantification of brain volumes.** To measure the volume of anatomical regions, the average RSTP reference brain (built using 40 STPT imaged brains) was registered to each brain sample, using the affine and b-spline registration procedure.[3,7] The number of voxels that belong to each region in the transformed STPT segmentation were counted and multiplied by 0.02 x 0.02 x 0.05 $mm^3$ (the dimensions of an anatomical voxel unit), resulting in the total volume of each region. Areas that passed the F.D.R. cutoff $q$<0.001 for contraction and $q$<0.05 for expansion were pseudo-colored (blue in case of contraction or red in case of expansion) for each dataset and the activation maps were overlaid on the average RSTP brain (e.g., Fig. 3).

**Seizure responses in 16p11.2 del/+ littermate.** The analysis included offspring of total 57 mutant del/+ mice from 25 breeder pairs, where each mutant had at least 1 mutant mate in the same litter. The probability of the same seizure response between litter 16p11.2 del/+ mice coming from the same litter was calculated using Poisson regression two models, with null hypothesis H0: mice have the same mean seizure score and H1: mice from the same father have the same mean seizure score (Supplementary Table 9).

**Homecage behavioral analysis and repeated seizure testing.** We analyzed homecage behavior of a cohort of 29 mice at 8 weeks of age. First, all mice were paired after weaning as 1 mutant and 1 wt male from the same litter per cage with water and food provided ad libitum. Second, at 8 weeks of age, homecage behavior was analyzed in four mice at a time by placing individual mice into four polycarbonate cages with dimensions of 48 × 26 × 20 cm (Ancare R20 series) during the light period, water and food were provided ad libitum on the wire bar of the ceiling of the cage. The behavior of each mouse was recorded and monitored with HomeCageScan (version 3.0; CleverSys) during 2 hrs before the beginning of the dark period and then 12 hours during the night. Each mouse was recorded on 4 different light/night days and was provided three days of rest in a homecage in between. Data were exported in bins of 5 min (24 bins 2hrs light, 100 binds first, second 6 hrs) and the selected 6 categories of behavior were analyzed by summing the time for each behavior for each day per mouse, then we calculated the mean values for the initial 2-hour period at light and the following two6-hour periods at dark. The behaviors included: 1) distanced traveled in meters; 2) fast movement: walk left, walk right, circle, forage, turn; 3) ceiling hanging: hang cuddled, hang vertically, hang vertically from rear up, hang vertically from hangs cuddled, land vertically, remain hang vertically, remain hang cuddled; 4) rearing category: come down, rear up, drinking, sniffing, remain rear up, remain partially reared, rear up from partially reared, rear up to partially reared; 5) grooming: repetitive licking and self-grooming behavior; 6) sleep/awake: sleep, twitch, awaken. After the homecage behavioral monitoring, all mice were tested three time for PTZ-evoked seizure response. The mice were injected three times, once per week, over a 3-week period with 30 mg/kg of PTZ and their behavioral response was video-recorded and scored off-line as described above.

**Bulk-RNA purification from blood and brain tissue in P2 pups (group 1)**. Pup mice were sacrificed on a postnatal day 2 before starting the dissection. We sterilized the instruments by heating them in a dry sterilizer or washing them with 70% (vol/vol) ethanol and drying it. We used 60-mm dishes with RNA protect solution tissue reagent during the dissection (Qiagen Cat No./ID: 76104). we euthanized the pups by decapitation and separated the head from the body. In parallel, we collected the blood for the pup's body (100-120 ul volume per animal). We employed tubes with a solution to protect the RNA from degradation for extended storage (Qiagen Cat No./ID: 76544). Place the head on a 60-mm dish and hold down the sides with forceps, gently dissect the skin on the top of the head and hold down the skin on either side with the forceps using a dissection microscope (stereo microscope). Using fine scissors, cut open the skull by making an incision at the base of the brain, separate the two halves of the skull and remove carefully. Take care not to cut through the brain tissue when removing the skull bone. It is essential to be extremely fast and careful to avoid contamination. Separate the two halves of the brain by making a sagittal cut along the midline. Discard the cerebellum. Place the brain such that the outer surface of the hemisphere faces the bottom of the dish. Under a dissecting microscope, gently remove the midbrain and thalamic tissue to leave an intact hemisphere containing the isocortex. Use another pair of forceps to pick and grab the meninges carefully and gently peel them off, ideally as a single piece. Check for the remaining parts of the meninges and remove them altogether. The isocortex from the right hemisphere immediately snapped frozen in liquid nitrogen for 30 seconds in cryotubes and then kept in dry ice to the final destination to the liquid-nitrogen tank. At P2, it is not possible to recognize the sex of the pups and were unknown at the moment of euthanization. For that, we collect a tail sample for each pup for DNA purification to detect the sex. It was determined using the Sry gene, and we used the male pups for posterior whole-genome expression sequence. The primers used SX_F, 5′-GATGATTTGAGTGGAAATGTGAGGTA-3′; SX_R, 5′-CTTATGTTTATAGGCATGCACCATGTA-3′. The Y chromosome is seen as a PCR amplicon band of 285 bp, and the X chromosome is detected when amplified a band of 685 bp (McFarlane L. et al., 2013).

**Bulk-RNA purification from brain tissue and blood from adult mice undisturbed (group 2).** The adult male mice were anesthetized with a Ketamine and Medetomidine mixture (60/0.5 mg/kg, IP). Then, we made terminal cardia blood withdrawal for the blood samples using a 1 ml TB syringe with a 25-29 gauge needle for blood collection (500-600 ul per mouse). The blood samples were kept in tubes with conservation solution for prolonged storage preservation (Qiagen Cat No./ID: 76554). They were preserved at -20 degrees until they proceeded to the RNA purification of all blood samples simultaneously. Then, the mice were decapitated and proceeded to peel off skin and muscles from the skull. Carefully, we peeled off the skull bone of the cerebellum part and made small cuts on the side by moving the scissors blades upwards to expose the cerebellum. Remove the skull very carefully by using the scissors as little as possible, cut carefully along the midline of the parietal by moving the scissors always away from the brain. Then, grab the parietal and remove it by bending it sideward and disconnecting the caudal part of the nasal and premaxilla from the frontal by carefully crunching the bone with forceps. After, the frontal along the midline caudally and rostrally grab the frontal edge and support the hold orbit. Twist gently and use one side of a forceps as the leverage to remove one part of the bone. Before taking the brain off from the skull, remove dura using forceps. We obtained a proximally 30 mg of brain tissue from the somatosensory area per adult mice, and we did fast snap freezing in liquid nitrogen using cryotubes. Then, we kept it in dry ice until delivery to the liquid-nitrogen tank.

**Brain biopsy surgery in adult mice (group 3).** The animals were induced by inhaling isoflurane (4 % to effect) using a small animal anesthetic induction chamber, surgical anesthesia, and monitoring breathing. The depth of anesthesia was monitored during the surgery. Animals were being placed on a heating pad on a small stereotaxic instrument. The body temperature was monitored and maintained at 37+-0.5 Celsius during the whole procedure. The whole procedure applied a layer of eye ointment to both eyes, putting hair removal lotion (Nair) on the mouse head, waiting for 1-2 minutes, and removing the hair and the lotion with cotton swabs. We disinfected the skin of the surgical region by applying betadine, 70 % alcohol, three times each. Then, we applied a thin layer of betadine to the skin. Inject 0.5 ml 5% glucose 0.9% saline mixed with atropine sulfate at concentration of 3 ug/ml subcutaneously, 40 ul 0.2% dexamethasone intramuscularly, meloxicam 5mg/kg subcutaneously, and 0.1 ml 0.2% Lidocaine +1:1000,000 Epinephrine in the surgical region subcutaneously. A paramedian incision at the right side of the midline was made by scalpel, and the skin on both sides of the incision was retracted with 25-0 silk sutures to expose the surgical field. Cotton swabs removed the periosteum on the dorsal part of the skull. The upper part (about 1 mm in width) of the temporal muscle was detached from the skull along the temporal ridge and cut off by a pair of spring scissors. The bleeding from the muscle was stopped by pressing gel foam or applying Vetbond glue on it. The craniotomy site was marked with the coordinates: AP +0.5 to – 4 mm and ML 1.5 to 6 mm from Bregma. The craniotomy was performed, and the bone flap was removed and put into sterilized saline in a petri-dish. The exposed cerebral cortex was cut (about 1 mm in depth) along the inner edge of the cranial window and was bluntly separated with the subcortical white matter by forceps. Once the separation was done, the removed brain tissue was quickly transferred into a precooled (by liquid nitrogen) tube, then put into dry ice until delivery in the liquid-nitrogen tank. The bleeding on the surgical field was stopped by pressing gel foam on it. The bone flap was repositioned at the cranial window site, sealed with Vetbond glue. The groove along the edge of the cranial window was filled with bone wax. The skin wound was closed by 6-0 absorbable sutures and covered by polysporin. The mice have injected Buprenorphine (0.075 mg/kg) and Baytril (10mg/kg) subcutaneously (SC), and it was put back into the home cage, and the recovery diet gel was used in the period of recovery. Mice were monitored 3 days after surgery. Baytril (10mg/kg) was injected once SC for 3 days after the surgery. Meloxicam (5mg/kg) was injected SC if the animals showed the signs of pain or distress three days after surgery.

**Seizure experiment after biopsy**. After animal surgery, we maintained each mouse in an isolated single cage for 3 days to recover. On day 4, the mice were injected with a single dose of 30 mg/kg PTZ convulsant drug injected intraperitoneally (i.p.) visually monitored by the experimenter for 30 min, and video-recorded for 90 min. After 90 min, the mice will be anesthetized with a 1.5x dose of ketamine (100 mg/kg) and dexmedetomidine hydrochloride (15 mg/kg) (i.p.) and euthanized to intracardiac blood extraction. The drug-evoked behavioral response was video-recorded, and the initial 30 min were manually scored off-line by two independent observers to quantify and classify the seizure-related behavior as we explained in the method sections "Quantification of PTZ-evoked responses". In mice, the PTZ compound shows behavior changes at 5 to 10' after i.p. injection.

**RNA purification and sequencing from brain tissue and blood samples.** We used TissueRuptor II (Cat. ID: 9002755-Qiagen) to prepare the homogenates from brain tissue of the day 2 postnatal animals and adult mice. Then, we used the RNeasy Plus Micro kit (Cat. ID: 74034-Qiagen) to purify the total RNA. We purified the RNA samples per age of all brain tissue samples simultaneously. For blood samples of P2 pups and adult mice, we used specific blood collections tubes to preserve the RNA (Cat. ID: 76544, 76544-Qiagen) and RNeasy Protect Animal Blood Kit (Cat. ID: 73224-Qiagen) to purify the total RNA, including the buffer RWT (Cat. ID: 1067933-Qiagen), to extract also microRNA. For quantification of RNA, we have used 1:10 dilution of the stock RNA and checked on Qubit using the Invitrogen Qubit™ RNA high sensitivity (HS) assay (Q33241). To analyze its quality, the Agilent RNA 6000 Nano kit (Cat. ID: 5067-1511). RNA integrity was assessed by the genomics facility prior to library preparation; all submitted RNA samples met facility acceptance criteria and were processed into libraries (median RIN = 8.6). Before preparing the libraries, we depleted globin for the blood samples using Kapa RNA HyperPrep Kit with RiboErase (KR1520-v2.17). For RNA-seq libraries preparation, we started with 250 ng of total RNA in 10 ul volume; we used multiplexed, pooled, and sequenced on multiple lanes of an Illumina NextSeq 500/550 Mid Output Kit v2.5 of 15M and 40M paired-end 150-cycles reads for each mouse sample.

**Methods to analyze the expression of the bulk-RNA for P2 and adult mice.** 100 gene signatures: To determine the molecular alterations present in 16p11.2 +/- mice, we conducted transcriptome analysis. Specifically, we performed RNA sequencing on brain samples collected from P2 pups and adult male mice (2-3 weeks old) and compared del/+ to their wt counterparts. FASTQ files were aligned to the GRCm39 mouse genome using STAR v2.7.4a7, and reads uniquely mapped to a gene were quantified using featureCounts (Rsubread v2.4.3). We compiled standard alignment QC metrics for each library from the processing pipeline, including total reads, mapping rate, uniquely mapped fraction, duplication rate, and rRNA fraction. QC metrics were reviewed within each age and tissue and did not differ systematically between comparison groups. Count normalization (median-of-ratios method), PCA, and differential expression testing were performed using DESeq2[9]. Principal components were clustered using *k*-means clustering with *k*=2. Fisher's exact test were used on the resulting clusters to determine separation of genotypes. Reported Log2-fold changes were shrunk using the ashr package.[10] All statistical analyses were performed using R (v4.03) in RStudio Server (v1.4). To construct and define the signatures, only the top 50% of highly expressed genes were used. After ranking the highly expressed genes, we selected the top 100 genes and aggregated them for genotype comparison. We normalized the wt groups to 1 and statistical analysis was performed using a t-test for two-group comparisons or one-way ANOVA for multiple-group comparisons using Bonferroni. The volcano plots include all genes, without cut-offs, so the bottom 50% of the genes (as ranked by DESeq2 baseMean expression level) are not found in the signatures. The IEGs expression signature contains 132 genes, and it was obtained from Wu et al. 2017 (Neuron Journal).[11] To address potential littermate effects, we performed a dedicated sensitivity analysis using cage identifiers in P2 and adult home cage cohorts. Because cage effects cannot be estimated reliably when partially confounded with genotype assignment, we restricted this analysis to mixed-cage subsets (cages containing both genotypes) and evaluated the predefined signature scores within these subsets. Genotype-associated signature effects remained robust within cage (adult brain: up-score $p = 0.0070$; down-score $p = 2.9\times10^{-5}$; P2 brain: up-score $p = 0.0087$; down-score $p = 1.25\times10^{-4}$), with consistent direction across all mixed cages. These results indicate that the key findings are not driven by cage structures.

**Functional enrichment:** To determine the functional relevance of genes with altered expression, functional enrichment was performed on the complete list of genes measured in the experiment, ranked by fold change using the Mann-Whitney U-test to evaluate differences in the distribution of genes associated with a GO term relative to all other genes (similar to PANTHER).[12] This analysis revealed an overrepresentation of genes associated with several neuronal GO terms among the genes upregulated in P2 and del/+ mice. This approach to functional enrichment enables the detection of subtle shifts in expression data that might be missed by methods requiring a pre-defined list of differentially expressed genes.

**Marker gene set enrichment:** Markers that robustly defining neuronal and non-neuronal cell types in the brain have been identified using data aggregated across multiple high-quality single-cell expression datasets, employing the combination of signatures published in 2021 to determine cell types.[13] While a change in the overall marker profiles would indicate compositional variation, the enrichment of cell type markers in the 100-gene signature would evince cell type-specific effects. We did not observe a change in the aggregate expression of the top 1000 markers for any neuronal or non-neuronal cell type in the del/+ vs. wt samples. Measuring the over-representation of such robust markers in the 100-gene signature defining the del/+ phenotype may shed some light on the cell types involved in driving this phenotype. Upon performing enrichment analysis using the hypergeometric test across the top 1000 markers for each cell type we discovered the overrepresentation of markers for multiple cell subclasses.[13] Evaluation of the overlaps of the 100 gene signature with each of the cell types against a co-expression network constructed by aggregating across multiple bulk datasets using EGAD showed that these common genes were strongly co-expressed.

**Statistical analyses.** Data points are stated and plotted as mean values ± SD, 95% CI, or box and whiskers (Tukey). $p$ values are represented by symbols using the following code: * for $0.01< p <0.05$, ** for $0.001< p<0.01$, and *** for $p <0.001$. Exact $p$-values are stated in figure legends and tables. For multiple comparisons statistical analysis, we employed the $q$ values. For mouse behavior experiments, each behavioral category was transformed to present an approximately normal distribution and in Figure 4 and Extended Figure 4a-b the statistical analysis were done using one-way ANOVA with Sidak's comparison test to compare two groups and one-way ANOVA and Tukey multiple comparison test to compare 3 groups.

**Statistical analysis of c-Fos$^+$ cell distributions.** Statistical comparisons between different PTZ concentrations and genotypes groups were done using either evenly spaced voxels or R.O.I.s based on the Allen Mouse Brain Atlas.[3,7] Voxels were overlapping 3D spheres with 150 µm diameter, spaced 20 µm apart; the cell count of each voxel was the number of GFP+ nuclei within 75 µm from the center of the voxel in 3D. We modeled the cell counts at a given location, $Y$, with a negative binomial distribution whose mean is linearly related to one or more experimental conditions, $X$: $E[Y]=\alpha+\beta X$. For example, when testing a saline solution versus a seizure group, our $X$ is a single column showing the categorical classification of mouse sample to group id, i.e. 0 for the control group and 1 for seizure group.[14,15] We found the maximum likelihood coefficients

$\alpha$ and $\beta$ through iteratively reweighted least squares, calculating the significance of the $\beta$ coefficient from estimates of its sample standard deviations. A significant $\beta$ means the group status is related to the cell count intensity at the specified location. The z-values in our summary tables correspond to this $\beta$ coefficient normalized by its sample standard deviation, which under the null hypothesis of no group effect, has an asymptotic standard normal distribution. The p-values give us the probability of obtaining a $\beta$ coefficient as extreme as the one observed by chance assuming this null hypothesis is true. To account for multiple comparisons across all voxel/R.O.I. locations, we thresholded the p-values and reported false discovery rates (F.D.R.) with the Benjamini-Hochberg procedure.[16] In contrast to correcting for type I error rates, this method controls the number of false positives among the tests that have been deemed significant. To compare voxel activation between the home cages and saline solution or saline solution and PTZ stimuli (e. g. Fig. 2, Supplementary Fig. 2), voxels that passed the F.D.R. cutoff $q<0.05$ were pseudo-colored (red) for each dataset and the activation maps were overlaid on the RSTP brain.

### Supplemental Figures

#### Supplementary Figure 1

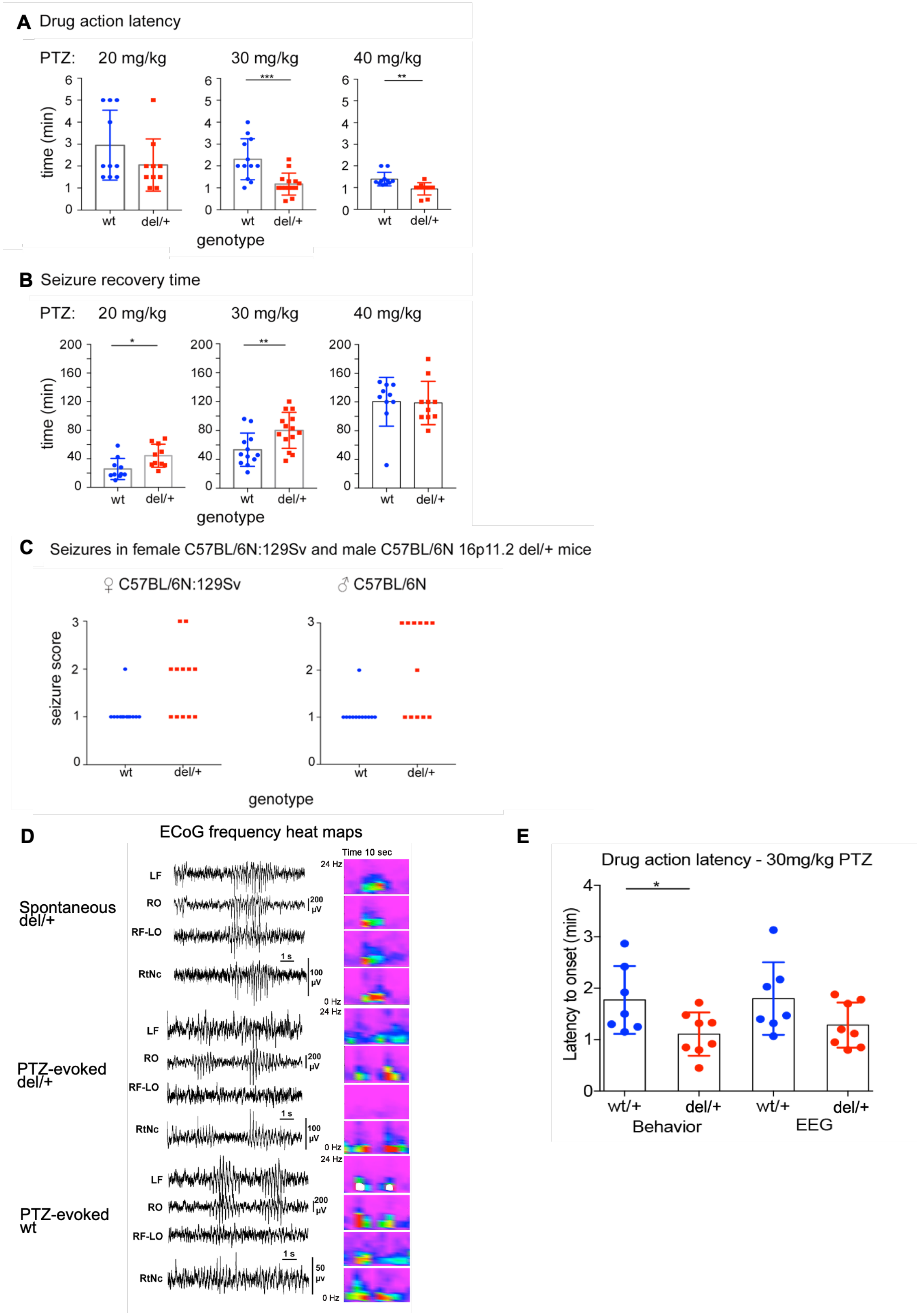

**Supplementary Figure 1 Seizure susceptibility in del/+ mice. (A)** Quantification of the latency (time to onset) of the first PTZ-evoked response in wt (blue) and del/+ (red) mice. No difference was seen at 20 mg/kg (left), but the latency was shorter in del/+ mice at both 30 mg/kg (middle) and 40 mg/kg PTZ (right): unpaired t-test $p<0.001$ at 30 mg/kg PTZ (wt *n*: 12, del/+ *n*: 14) and $p<0.01$ at 40 mg/kg PTZ (wt *n*: 11, del/+ *n*: 11 mice). **(B)** Quantification of the recovery time, defined as the time of first crossing the length of the cage after PTZ-evoked effect[6] in wt (blue) and del/+ (red) mice. The recovery time was longer in del/+ mice at PTZ 20 mg/kg (left) and 30 mg/kg (middle): unpaired t-test $p<0.05$ for 20mg/kg (wt *n*: 10 and del/+ *n*: 10) and $p<0.01$ for 30 mg/kg (wt *n:* 12 and del/+ *n*: 14). **(C)** Seizure susceptibility in female F1 C57BL/6N:129Sv and male 100% C57BL/6 background del/+ (red) mice. 30 mg/kg PTZ evoked varied responses in both groups of del/+ mice: 5 out 12 myoclonic seizures and 2 out of 12 clonic seizures in female del/+ (C57BL/6N:129Sv) mice, and 1 out 12 myoclonic seizures and 6 out 12 clonic seizures in male del/+ (100% C57BL/6). In contrast, only freezing was seen in all but 1 wt (blue) mice of both groups (wt female *n*: 13 and wt male *n*: 12). The seizure severity score is as in Figure 1A. **(D)** Frequency heat maps of ECoG recordings shown in Figure 1E-G demonstrating both cortical and reticular thalamus circuit participation in spontaneous and PTZ-evoked synchronous events. **(E)** Shorter latency of PTZ-evoked behavioral response in del/+ mice with ECoG recordings. The time to onset to the first PTZ-evoked behavioral (behavior) and ECoG seizure response (ECoG) in wt (blue) and del/+ mice (red). Statistical comparison by Mann-Whitney test (* for $p < 0.05$).

### Supplementary Figure 2

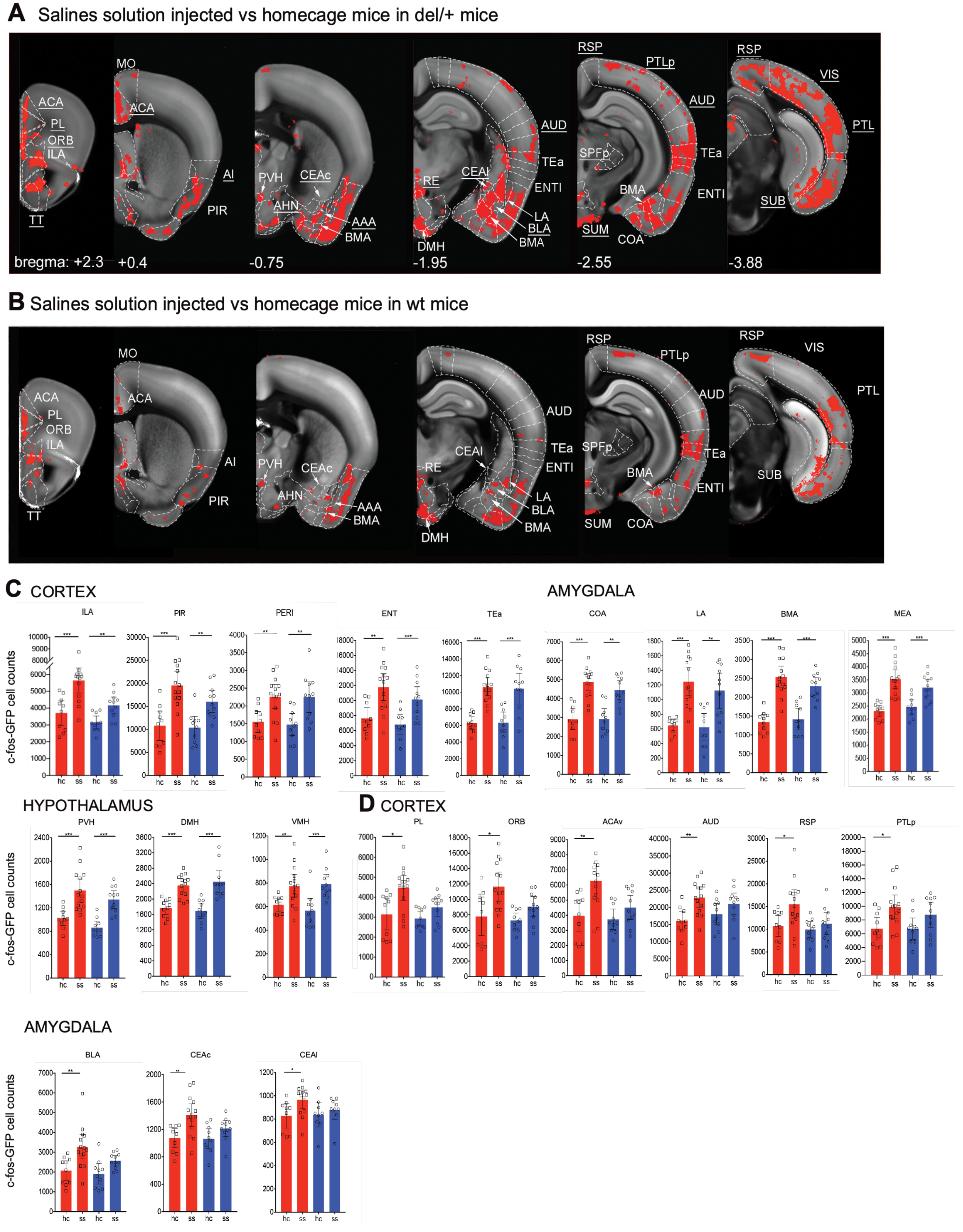

**Supplementary Figure 2 Quantification of saline i.p. injection-evoked neuronal activation by whole-brain c-Fos$^+$-mapping points to analyze the excitability in del/+ mice.** Statistical comparison of c-Fos$^+$ cell distribution between undisturbed mice (homecage group) and mice injected i.p. with saline (control saline group), with significant increases representing higher brain activation in the saline group shown as red areas overlaid on a grayscale RSTP mouse brain[3,17] (Methods). **(A-B)** The moderate stress of i.p. saline injection evoked comparable activation of stress-related structures in del/+ and wt mice, including infralimbic area (ILA), piriform cortex (PIR), paraventricular hypothalamic nucleus (PVH), lateral amygdalar nucleus (LA), basomedial amygdalar nucleus (BMA), dorsomedial of the hypothalamus (DMH), temporal association areas (TEa), entorhinal area, lateral part (ENTl), cortical amygdalar area (COA). In addition, several structures were activated only in del/+ mice (abbreviations underlines in A), including the following cortical areas: anterior cingulate (ACA), prelimbic (PL), orbital (ORB), and infralimbic (ILA), agranular insular (AI), auditory (AUD), retrosplenial (RSP), posterior parietal association (PTLp) and visual (VIS) cortex, as well as several subcortical areas: taenia tecta (TT), basolateral amygdalar nucleus (BLA), central amygdalar nucleus, capsular part (CEAc), central amygdalar nucleus, lateral part (CEAl), anterior amygdalar area (AAA), anterior hypothalamic nucleus (AHN), supramammillary nucleus (SUM), and subiculum (SUB) (wt homecage *n*: 10, wt saline injected *n*: 11, del/+ homecage *n*: 10, del/+ saline injected *n*: 14). The statistical *q* values were derived by negative binomial regression corrected for multiple comparisons by false discovery rate (FDR) $*q<0.05$. **(C)** Bar graph representation of c-Fos$^+$ cell distribution in selected brain areas with similar c-fos induction in del/+ (red) and wt (blue) mice. Additional abbreviations not listed in A-B are: perirhinal area (PERI), medial amygdalar nucleus (MEA), ventromedial hypothalamus nucleus (VMH). **(D)** Bar graph representation of c-Fos$^+$ cell distribution in selected brain areas with higher c-$Fos^+$ induction in del/+ (red) than wt (blue) mice. The statistical *q* values were derived by negative binomial regression corrected for multiple comparisons by false discovery rate (FDR); *$q<0.05$, **$q<0.01$, ***$q<0.001$. hc (homecage mice) and ss (saline solution injected mice).

#### Supplementary Figure 3

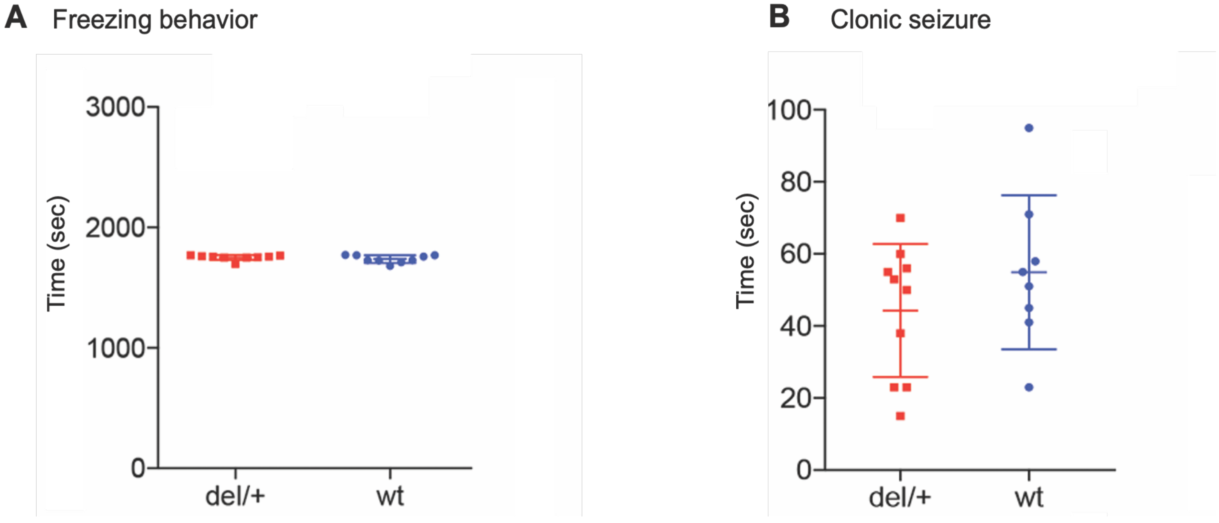

**Supplementary Figure 3 Quantification of PTZ-evoked seizures with 30 mg/kg for del/+ and 35 mg/kg for wt mice. (A)** Quantification of the duration of freezing behavior evoked by PTZ 30 mg/kg in del/+ (red) and 35 mg/kg in wt (blue) mice. These animals were used for the comparison in Fig 2A, and C. Unpaired t-test did not show significant differences. **(B)** Quantification of the duration of clonic seizure evoked by PTZ 30 mg/kg in del/+ (red) and 35 mg/kg in wt (blue) mice. These animals were used for the comparison in Fig 2B, and 2D. Unpaired t-test did not show significant differences.

#### Supplementary Figure 4

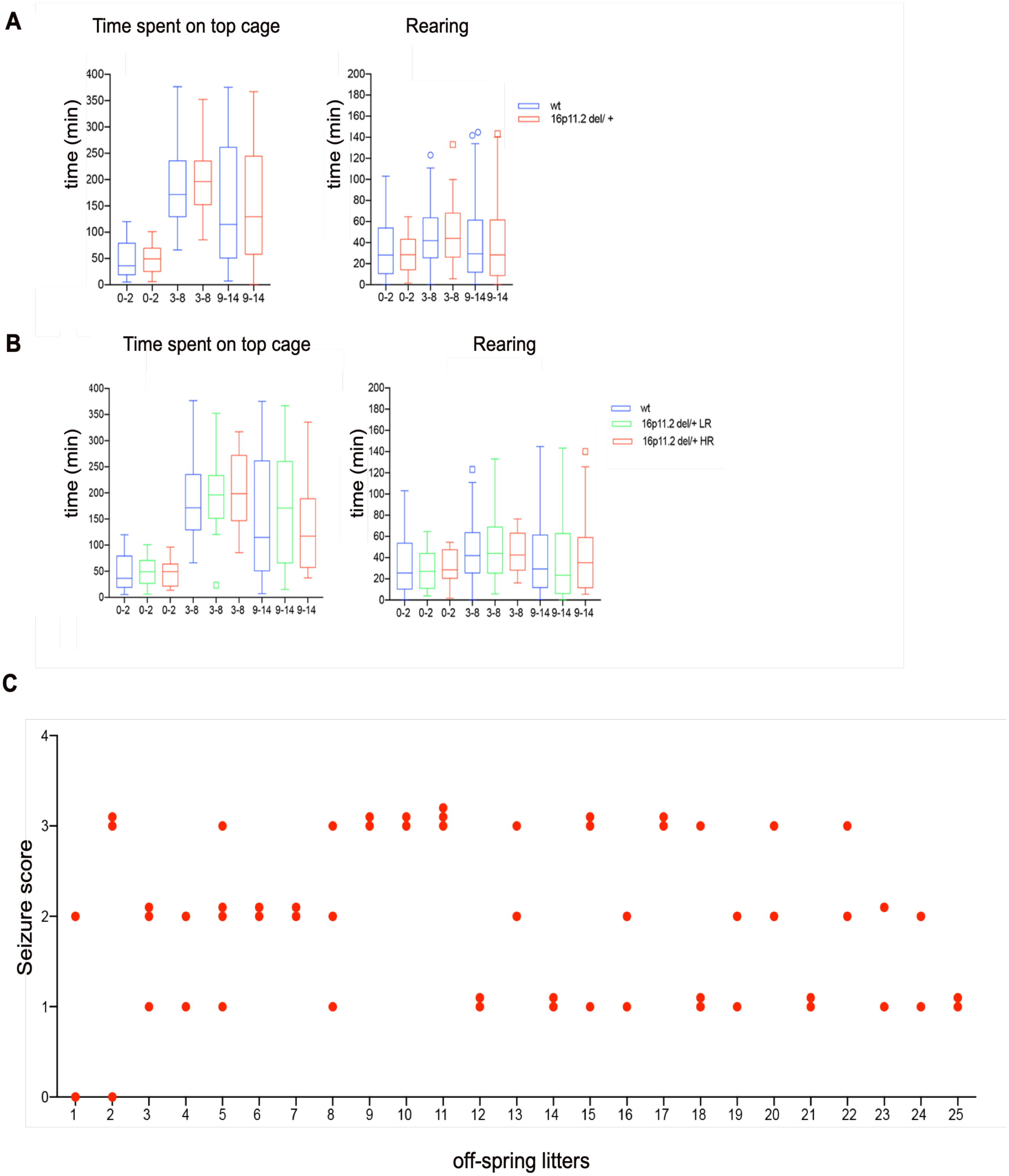

**Supplementary Figure 4 Additional analyses of seizure within 16p11.2 litters. (A)** Two-cohort comparison between all del/+ (red) mice (*n* = 17) and wt (blue) littermates (*n* = 12) showed no group differences for the time spent on the top of the cage and time spent in rearing. **(B)** Three-cohort comparison between del/+ LR mice (green) (*n* = 9), del/+ HR mice (red) (*n* = 8), and wt littermates (blue) (*n* = 12) showed also no group differences for the time spent on the top of the cage and time spent in rearing. **(C)** Seizure scores of del/+ mice within the same littermates. The analysis of PTZ 30 mg/kg evoked responses in del/+ mice from 25 litters with 2 or more del/+ siblings by Poisson regression showed no correlation between direct siblings and seizure score ($p$ = 0.97). Each dot represents one del/+ mouse.

#### Supplementary Figure 5

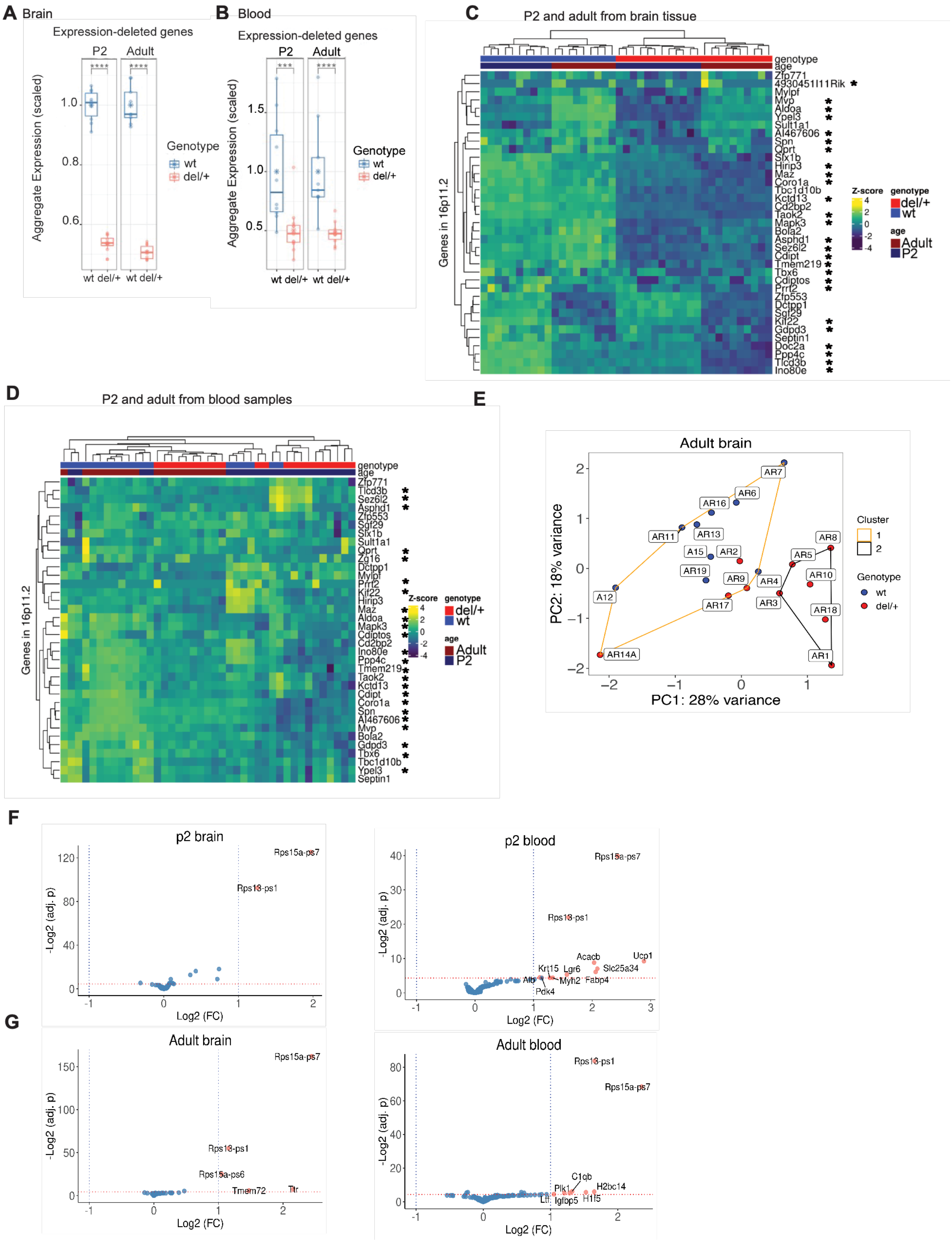

**Supplementary Figure 5 Bulk RNA purification from tissue and blood samples confirms the heterozygosity of the genes in the deletion in P2 and adult del/+ animals. (A)** Aggregated expression to analyze the genes in the deletion of del/+ (red) from P2 and adult mice for brain tissue (groups 1 and 2) (**** for $p$ <0.0001) (**** for $p$ <0.0001). (**B)** Blood (*** for $p$ <0.001) (**** for $p$ <0.0001). (**C)** Heat map, from the genes included in the deletion from the del/+ mice deletion in P2 animals and adult homecage. The differential expression list for the 26 genes in the 16p11.2 deletion. The heat map also includes genes immediately outside of the deletion. (*) Genes included in the deletion. **(D)** Heat-map including the genes in the del/+ mice in P2 and adult mice from blood samples. The heat map also includes genes immediately outside of the deletion. (*) Genes included in the deletion. **(E)** Principal component analysis of adult brain tissue including the information of each mouse in cluster 1 and 2 to show the number of mice per cluster. (**F)** Volcano graphs showing the differential gene expression vs. p values in the P2 animals in brain tissue and blood (adj. $p$<0.05). Upregulates genes (blue) and downregulated genes (blue) **(G)** Volcano graphs showing the differential gene expression vs. p values in the adult group in brain tissue and blood (adj. $p$<0.05). Upregulates genes (blue) and downregulated genes (blue).

#### Supplementary Figure 6

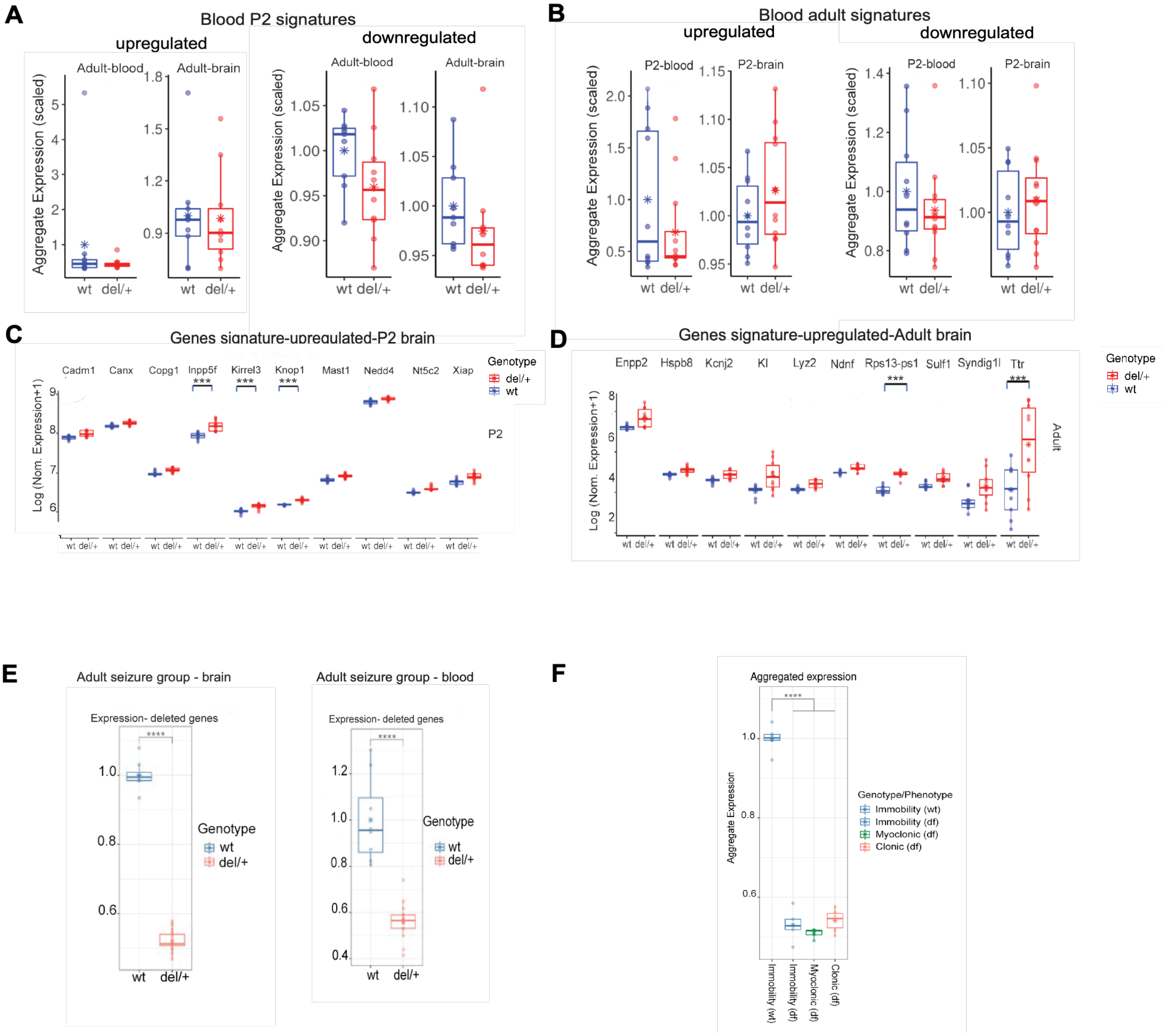

#### Supplementary Figure 6

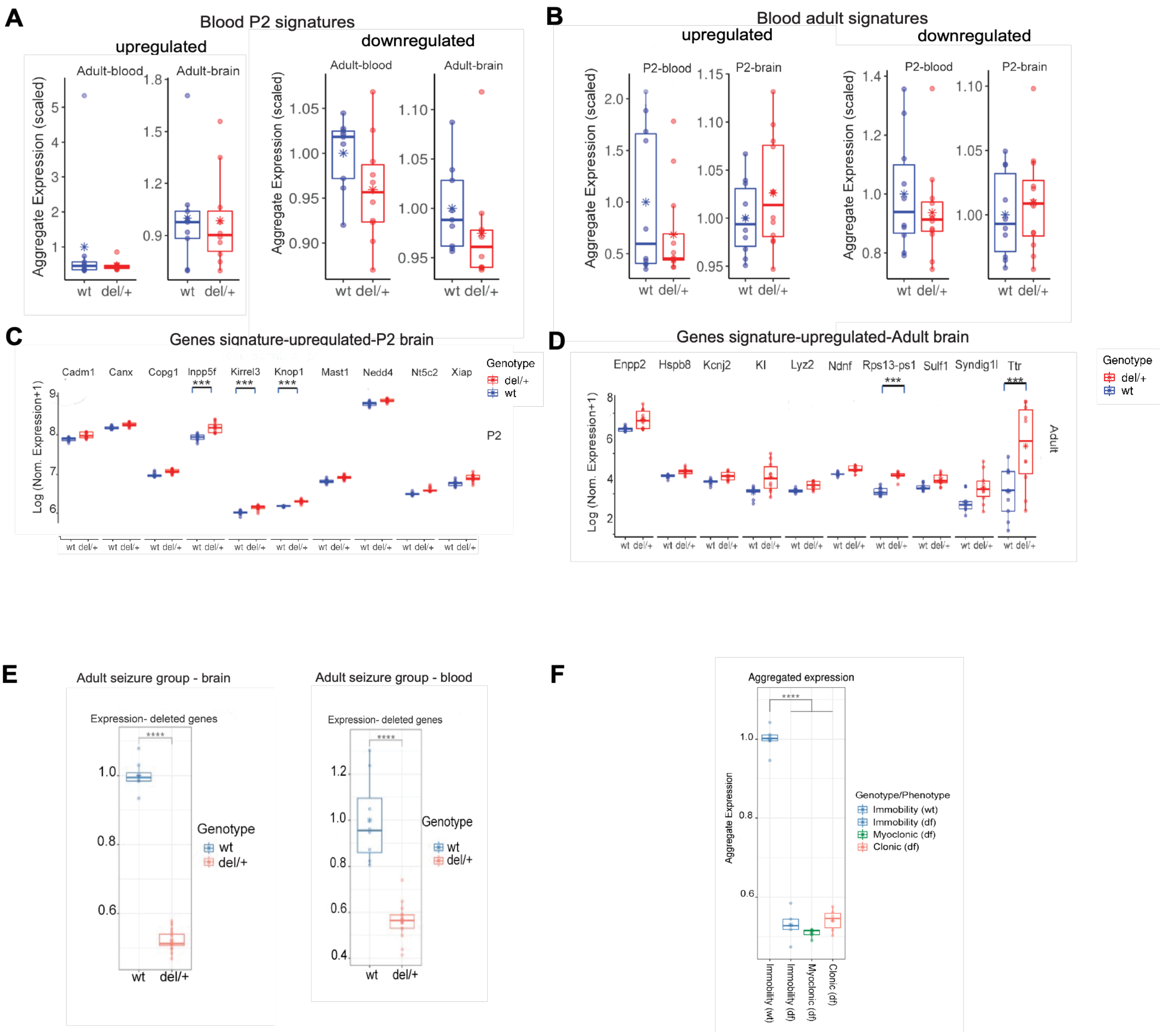

**Supplementary Figure 6 Differences in gene expression between P2, adult, and PTZ-treated adult. (A)** Top 100 upregulated and downregulated gene signatures from P2 blood in blood samples and brain tissue of adult mice homecage. Del/+ (red), wt (blue) **(B)** Top 100 upregulated and downregulated gene signatures from adult blood in blood samples and brain tissue of P2 mice. These signatures were not useful to be applied in different ages and samples for genotype differentiation. Del/+ (red), wt (blue) (graphs indicate mean expression and SD-/+). (**C)** Top 10 genes from the 100 upregulated genes of P2 animals including the genes that were statistically significant in the differential expression analysis (*lnpp5f, Kirrel3 and Knop1*). (**D**) Top 10 genes from the 100 upregulated genes of adult animals and the genes that were statistically significant in the deferential expression analysis (*Rsp13-ps1*, *Ttr,* significant differences). **(E)** Aggregated expression to analyze the genes in the deletion of del/+ mice in the seizure animals and their wt littermates (group 3) (**** for $p$ <0.0001) (**** for $p$ <0.0001). (**F)** Aggregated expression to analyze the genes in the deletion distinguished by seizure level (**** for $p$ <0.0001).

#### Supplementary Figure 7

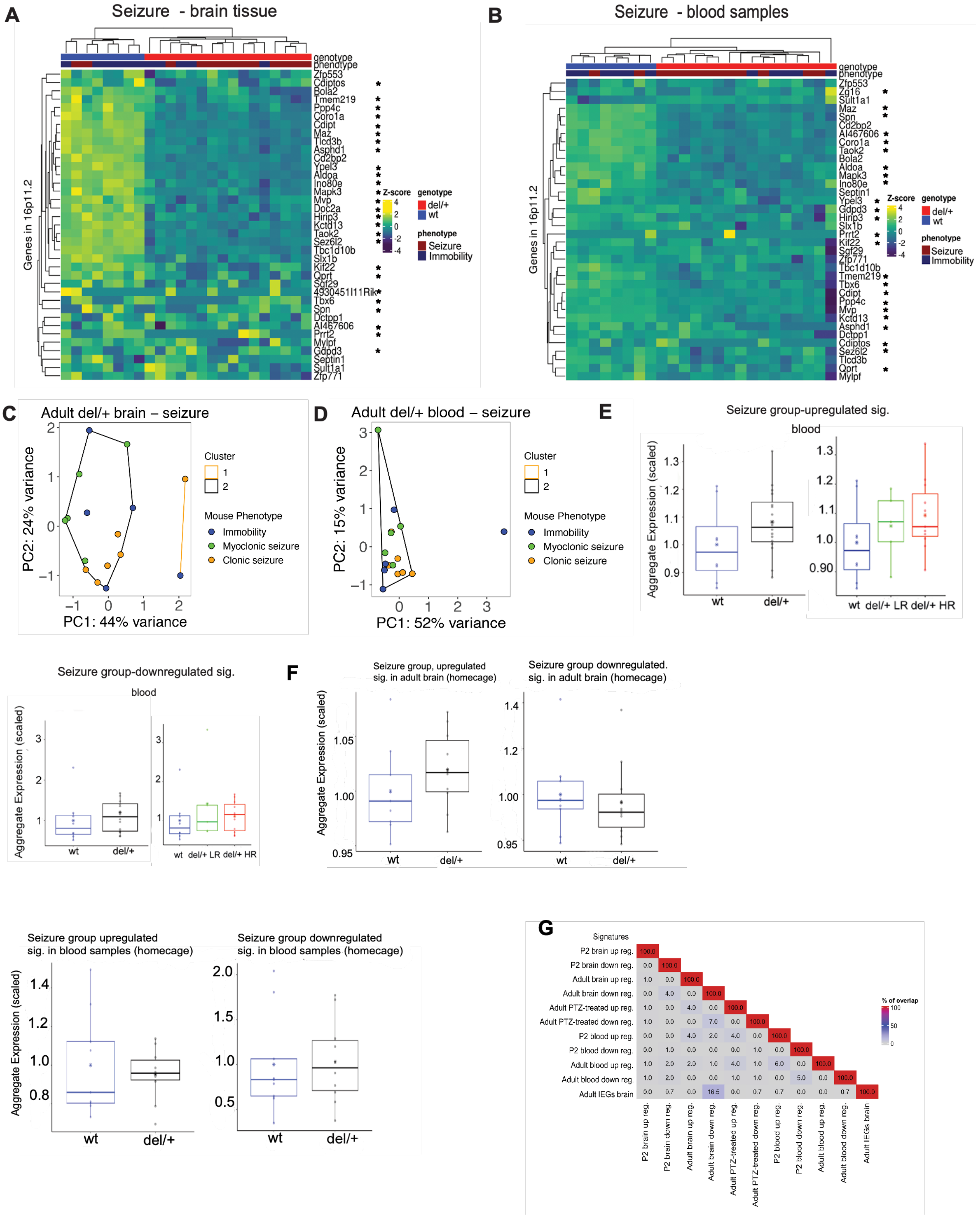

**Supplementary Figure 7 Analysis of the adult signatures from the home cage group and the PTZ-treated group. (A)** Heat map including the genes in the deletion in the PTZ-treated group mice from the brain (group 3) (*) Genes included in the deletion. **(B)** Heat map including the genes in the deletion in the PTZ-treated group mice from blood samples (group 3) (*) Genes included in the deletion. **(C)** Principal component analysis in the del/+ adult mice treated with PTZ using the mouse seizure levels, which still cannot separate the different clusters using the RNA from brain tissue. **(D)** Principal component analysis in the del/+ adult mice treated with PTZ which still cannot separate the different clusters using the RNA from blood. **(E)** Top 100 upregulated and downregulated gene signatures from the PTZ-treated adult group (group 3) using blood samples. These signatures were not useful for genotype separation. Del/+ (black), wt (blue), LR (green) and HR (red) **(F)** Top 100 upregulated and downregulated genes signatures from the PTZ-treated adult mice from brain tissue applied in the adult group (homecage). Top 100 upregulated and downregulated gene signatures from the PTZ-treated adult mice applied to the adult group from blood samples (homecage)**.** (**G)** Comparison of all signatures to analyze the percentage of overlap between them from brain tissue and blood samples.

### Supplementary Figure 8

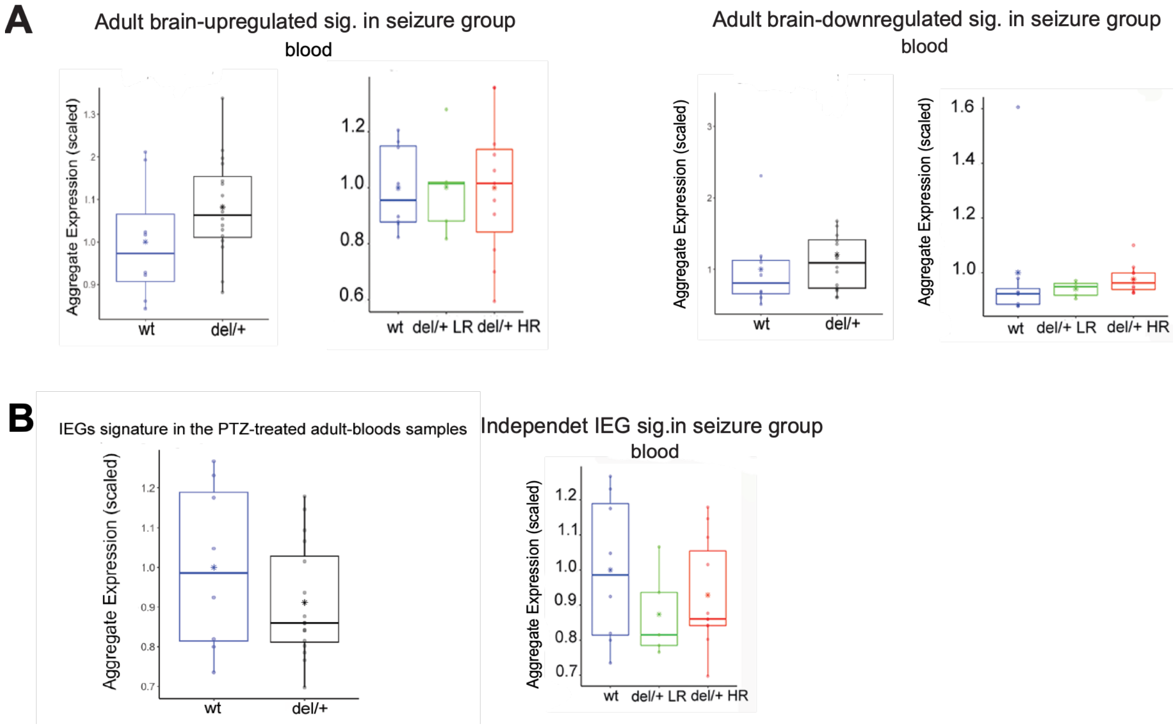

**Supplementary Figure 8 (A)** Top 100 upregulated and downregulated gene signatures from adult brain group 2 (homecage) applied to blood samples from PTZ-treated adults. Del/+ (black), wt (blue), LR (green) and HR (red) **(B)** The IEGs signature was applied to blood samples from the PTZ-treated adult group. The blood signatures are not useful for del/+ vs. wt genotype separation. Del/+ (black), wt (blue), LR (green) and HR (red)

**Supplemental Table Legends**

**Supplementary Table 1 Statistical analysis of c-Fos$^+$ distribution in undisturbed (homecage) and saline injected (saline control) del/+ and wt mice related to Supplementary Figure 2.** (**A)** A comparison between undisturbed homecage del/+ and wt mice revealed largely the same c-fos+ cell distribution, suggesting a lack of gross difference in baseline neural activity at rest. (**B)** A comparison between wt homecage and saline-injected wt mice revealed modest brain activation of mainly stress-related structures that are color-coded based on statistical significance (see right side of the data columns). (**C**) A comparison between del/+ homecage and saline-injected del/+ mice revealed similar modest brain activation of stress-related structures and in addition activation of several cortical areas not seen activated in wt mice under the same conditions. Such activation agrees with the increased cortical excitability evoked by PTZ, as shown in Figure 1 and 2. The activated structures are color-coded based on statistical significance. Columns in each excel sheet represent: Column A = full anatomical names of the brain structures, B = abbreviation of the anatomical names, C = hierarchical structure order, D-G = mean and SD of c-fos-GFP+ cell counts of the groups being compared, H = % change between the groups being compared, I-K = the statistical z-scores, uncorrected p values, and corrected FDR q value for the comparison between the groups being compared. (**D**) Side by side comparison between saline injection evoked c-fos+ cell increase in wt mice (left) and del/+ mice (right), with color-coded statistical significance as in done in Supplementary Table 1A-C.

**Supplementary Table 2 Statistical analysis of c-Fos$^+$ distribution in del/+ saline control and 30 mg/kg PTZ-injected mice and in wt saline control and 35 mg/kg PTZ-injected mice related to Figure 2.** (**A**) A comparison between del/+ saline control and LR del/+ score 1 PTZ-evoked response revealed modest PTZ-evoked activation that, however included several structures activated only in the del/+ score 1 mice. Those structure are highlighted in yellow in column L. (**B**) A comparison between del/+ saline control and HR del/+ score 3 PTZ-evoked response revealed broad PTZ-evoked activation that included many cortical and subcortical areas. **(C)** A comparison between wt saline control and wt score 1 PTZ-evoked response revealed modest PTZ-evoked activation of mainly subcortical areas. (**D**) A comparison between wt saline control and wt score 3 PTZ-evoked response revealed broad PTZ-evoked activation that includes many cortical and subcortical areas. The activated structures are color-coded based on statistical significance. Columns in each sheet represent: Column A = full anatomical names of the brain structures, B = abbreviation of the anatomical names, C = hierarchical structure order, D-G = mean and SD of c-fos-GFP+ cell counts of the groups being compared, H = % change between the groups being compared, I-K = the statistical z-scores, uncorrected p values, and corrected FDR q value for the comparison between homecage and saline group c-Fos+ cell distributions. **(E)** Side by side comparison between brain activation responses of: LR del/+ mice with L1 seizure (columns B-C), and HR del/+ mice with L3 seizure (columns D-E), wt mice with L1 seizure (columns F-G), and wt mice with L3 seizure (columns H-I), with color-coded statistical significance as in done in (**A-D)** In addition, yellow color in column A marks the ROIs with significant activation seen only in the LR del/+ mice with L1 seizure. Raw data for Table 2A, Raw data for Table 2B, Raw data for Table 2C and Raw data for Table 2D. This data contains the ROI anatomical areas and the number of positive c-Fos+ cells per animal.

**Supplementary Table 3 Statistical analysis of brain volume changes in wt and del/+ baseline, LR and HR mice related Figure 3. (A)** A brain volume comparison between naive (pooled homecage and saline injected) wt and del/+ mice revealed that del/+ displayed volume reductions of the caudoputamen, hippocampus, and several cortical areas, including the frontal, motor, somatosensory, auditory, and visual cortex, and volume enlargements of the hypothalamus, superior colliculus, and periaqueductal gray. The data are color-coded based on statistical significance (see right side of the data columns). (**B**) A brain volume comparison between wt mice with PTZ-evoked seizure score 1 and LR del/+ mice with PTZ-evoked seizure score 1 revealed subcortical expansions in LR del/+ mice seen in the earlier naïve mouse comparison, including the periaqueductal gray and superior colliculus, but only minor reductions in the cortex and elsewhere. (**C**) A brain volume comparison between wt mice PTZ-evoked seizure score 3 mice and HR de/+ mice with PTZ-evoked seizure score 3 revealed large brain volume reductions in HR del/+ mice, including all cortical areas, caudoputamen, hippocampus and many thalamic nuclei, but without any subcortical enlargements. Columns in each excel sheet represent: Column A = full anatomical names of the brain structures, B = abbreviation of the anatomical names, C = hierarchical structure order, D-G = mean and SD of c-fos-GFP+ cell counts of the groups being compared, H = % change between the groups being compared, I-K = the statistical z-scores, uncorrected p values, and corrected FDR q value for the comparison between the groups being compared. (**D**) Side by side comparison of brain volumes for naïve (pooled) del/+ mice (A-B), LR del/+ mice (C-D) and HR del/+ mice (E-F). Raw data of the volume in pixels per animals from homecage and saline injected from Table 3A. Raw data of the volume in pixels per animals from wt score 1 and 16p11.2 del/+ LR score 1 from Table 3B. Raw data of the volume in pixels per animals from wt score 3 and 16p11.2 del/+ HR score 3 from Table 3C. Volume comparison between 16p11.2 del/+ LR vs 16p11.2 del/+ HR with $q<0.05$.

**Supplementary Table 4 PCA analysis for genotype separation between del/+ and wt for group 1, group 2, and group 3 for Figure 5 and Figure 6.** Brain-P2 (group 1), brain-adults (group 2), brain -adult with seizure (group 3), brain- adult del/+ mice only (group 3), blood- P2 mice (group 1), blood-adult mice (group 2), blood-adult with seizure (group 3), blood- adult del/+ mice only (group 3).

**Supplementary Table 5 List of top 100 genes of all the signatures employed in the analysis for Figure 5 and Figure 6 from brain and blood samples.** List of 100 top upregulates and downregulated genes used for all the groups, including *log2FoldChange*, *p-value*, and *an adjusted p-value* for brain and blood compared with the wt mice. Constructed signatures from brain tissue: (**A)** P2 up brain: P2 pups upregulated genes. (**B)** P2 down brain: P2 pups downregulated genes. (**C)** Adults up brain: Adult upregulated genes. (**D)** Adult down brain: Adult downregulated genes. (**E**) Seizure up brain: Adult seizure group upregulated genes. **(F)** Seizure downregulated brain: Adult seizure group downregulated genes. Constructed signatures from blood: (**G)** P2 up blood: P2 pups upregulated genes. (**H)** P2 down blood: P2 pups downregulated genes. (**I)** Adults up blood: Adult upregulated genes. (**J)** Adult down blood: Adult downregulated genes. (**K)** Seizure up blood: Adult seizure group upregulated genes. (**L)** Seizure downregulated blood: Adult seizure group downregulated genes. **(M)** LR (low responder) and HR (high responder) upregulated and downregulated signatures.

**Supplementary Table 6 List of the 10 top upregulated genes from the P2 and adults for Supplementary Figure 6C-D. (A)** List the 10 top genes from the 100 top upregulated genes from P2 pups and their adjective p-value in adult genes. (**B)** List the 10 top genes from the 100 top upregulated genes signature in adults and their adjective p-value in P2 pups. (**C)** The rank of the 10 top upregulated genes from the adults in the whole-transcriptomic RNA in the P2 pup.

**Supplementary Table 7 Lists of the GO enrichment pathways, including the genes involved that are affected in brain tissue and blood samples for the Figure 5 and Figure 6. (A)** For P2 (group 1), adult home cage (group 2), seizure group (group 3), del/+ LR and del/+ HR up and downregulated signature genes in brain tissue with the adjusted p-value information. **(B)** For P2 (group 1), adult home cage (group2), seizure group (group 3), del/+ LR and del/+ HR up and downregulated signature genes in blood with the adjusted p-value information.

**Supplementary Table 8 Statistical analysis of the seizure group in the Figure 6 C, D, E, F. for brain and blood. (A)** Adjustive p-values results from the seizure group (del/+ LR and del/+ HR) 100 top upregulated genes from brain and blood. **(B)** Adjustive p-values results from seizure group (del/+ LR and del/+ HR) 100 top downregulated genes from brain and blood. **(C)** Adjustive p-values from adult 100 top upregulated genes signature applied in the seizure group (del/+ LR and del/+ HR) in brain and blood. **(D)** Adjustive p-values from adult 100 top upregulated genes signature applied in the seizure group (del/+ LR and del/+ HR) in brain and blood. **(E)** Adjustive p-values from I.E.G.s signature applied in the seizure group (del/+ LR and del/+ HR) in brain and blood.

**Supplementary Table 9 Parental and seizure information for all cases with 2 or more 16p11.2 mice born within a litter related to Figure 1 and Discussion.** 25 such litters were analyzed (column A), from 21 breeding cages (column B), with PTZ-evoked seizure responses (column E), scored based on the behavioral response as shown in Figure 1: immobility (score 1), myoclonic seizure (score 2), clonic seizure (score 3), tonic-clonic seizure (score 4) (column F), and genotype of the animals (column G).
